# Evolutionary analysis reveals the role of a non-catalytic domain of peptidyl arginine deiminase 2 in transcriptional regulation

**DOI:** 10.1101/2022.09.19.508513

**Authors:** José Luis Villanueva-Cañas, Narcis Fernandez-Fuentes, Dominik Saul, Robyn Laura Kosinsky, Catherine Teyssier, Malgorzata Ewa Rogalska, Ferran Pegenaute Pérez, Baldomero Oliva, Cedric Notredame, Miguel Beato, Priyanka Sharma

## Abstract

Peptidyl arginine deiminases (PADIs) catalyze protein citrullination, a post-translational conversion of arginine to citrulline. The most widely expressed member of this family, PADI2, regulates cellular processes that impact several diseases. We hypothesized that we could gain new insights into PADI2 function through a systematic evolutionary and structural analysis. Here, we identify 20 positively selected PADI2 residues, 16 of which are structurally exposed and maintain PADI2 interactions with cognate proteins. Many of these selected residues reside in non-catalytic regions of PADI2. We validate the importance of a prominent loop in the middle domain that encompasses PADI2 L162, a residue under positive selection. This site is essential for interaction with the transcription elongation factor (P-TEFb) and mediates active transcription of the oncogenes *c-MYC*, and *CCNB1,* as well as impacting cellular proliferation. These insights could be key to understanding and addressing the role of the PADI2 *c-MYC* axis in cancer progression.

**Significance Statement:** Here we use a systematic evolutionary analysis to identify positively selected residues in the non-catalytic domain of PADI2 and link the positive selection of key residues to a role in transcription. Specifically, a structurally exposed loop in the PADI2 middle domain encompasses the positively selected residue L162 which is linked to transcription and cellular proliferation. This loop contributes to PADI2 interaction with the P-TEFb complex and cellular proliferation. Our results showcase the utility of combining evolutionary and experimental approaches to dissect the evolution of essential functional processes.

## Introduction

Members of the peptidyl arginine deiminase (PADI) family catalyze post-translational calcium-dependent citrullination of arginines, converting them into the non-coded amino acid citrulline [1–4]. Arginine citrullination is a widespread post-translational modification (PTM) that increases the hydrophobicity of proteins and can contribute to fine-tuning physiological processes. For instance, citrullination of core histones weakens histone–nucleic acid interactions, impacting both chromatin organization and transcription [5]. Citrullination can also affect protein folding and consequently protein function [6–9]. The five members of the PADI family have distinct tissue-specific expression patterns: PADI1 is present only in the epidermis and uterus; PADI3, in the epidermis and hair follicles; PADI4, in immune cells, brain, uterus, and bone marrow; PADI6, in ovarian egg cells, embryonic tissues, and testicles; and PADI2, in brain, uterus, spleen, breast, pancreas, skin, and skeletal muscles [2,3]. Notably, PADI2 has been associated with multiple pathological states, including autoimmune disorders [10] and neurological diseases [11,12], as well as with several cancers (e.g., breast, cervical, liver, lung, ovarian, thyroid, gastric, and prostate cancer) [13–22]. The link between PADI2 and various diseases underscores an unmet need to elucidate its molecular mechanisms of action and to understand its pathophysiological function.

PADI family members have distinct substrates and target diverse arginine residues in different proteins [23]. All family members are Ca^2+^-dependent, but PADI2 also relies on ordered calcium binding to its active site for substrate binding and catalysis, as demonstrated in the deimination of arginine 26 in histone H3 (H3R26) [24–27]. Importantly, H3R26 citrullination is critical for interaction with the chromatin remodeller SMARCAD1 (SWI/SNF-related matrix-associated actin-dependent regulator of chromatin subfamily A containing DEAD/H box 1) and hence links PADI2 to gene activation in the context of naive pluripotency [28]. PADI2 also contributes to chromatin modification that promotes the differentiation of oligodendrocyte precursors and efficient myelination, processes required for motor and cognitive functions [29]. Recently, PADI2 was found to citrullinate MEK1, thereby promoting signaling by extracellular signal-regulated protein kinase 1/2 (ERK1/2) signaling in endometrium cancer [22].

Recently, we identified another unique function of PADI2: the deimination of arginine 1810 (R1810) on repeat 31 of the C-terminal domain (CTD) of the large subunit of RNA polymerase 2 (RNAPII). This modification potentiates the interaction between the RNAPII-CTD and the positive transcription elongation factor b complex (P-TEFb) and consequently facilitates gene expression that is essential for cell identity [30]. Indeed, we reported that citrullination of R1810 in the CTD of RNAPII facilitates its interaction with the P-TEFb kinase complex and promotes the recruitment of CDK9 to transcription start sites (TSS) [30]. This contributes to overcoming the RNAPII pausing barrier, highlighting the functional connection between PADI2 and the P-TEFb kinase complex. These studies support the notion that PADI2 can selectively citrullinate arginine residues in specific proteins, which may explain its relevance to several pathophysiological conditions.

The *PADI* genes are ubiquitous in vertebrates but are absent from yeast, worms, and flies. A recent study focusing on the comprehensive identification of *PADI* homologs unveiled the evolutionary trajectory of *PADIs* within the animal lineage [31]. *PADIs* appear to have been introduced from cyanobacteria into animals by horizontal gene transfer (HGT) [31], supporting the previous hypothesis of HGT as a mechanism for introducing new genetic material into vertebrate genomes [32–35]. Enzymes mediating citrullination in human parasites and microbes are highly divergent in sequence and have different substrate specificities. These include pPAD, which is an extended agmatine deiminase found in *Porphyromonas gingivalis*, and *Giardia lamblia* ADI, an extended form of the free L-arginine deiminase gADI found in this human parasite [36,37]. Previous small-scale phylogenetic analyses of the PADI family have shown that PADI2 is the most conserved family member. Mammalian PADIs comprise three structural domains; the N-terminal domain (NTD) (PADI_N, Pfam annotation: PF08526), the middle domain (PADI_M, Pfam annotation: PF08527), and the catalytic C-terminal domain (PADI_C, Pfam annotation: PF03068) [38–41]. Of note, PADI1 and PADI3, and PADI4 and PADI6, are more closely related in evolutionary terms than PADI2 and any other family member [6]. This finding suggests that PADI2 is most closely related to the ancestral PADI enzyme and that the other PADI enzymes arose through gene duplication more recently in evolution.

The enzymes responsible for essential PTMs (including phosphorylation, acetylation, and glycosylation) are found across all domains of life, suggesting that they were present in the Last Universal Common Ancestor (LUCA) [42,43]. Similarly, the PADI_C domain was present in the LUCA, indicating an ancient origin of citrullination. The PADI_M domain, which encompasses the sequential calcium-binding sites and maintains allosteric communication with PADI_C [25], is present in cyanobacteria. Indeed, recent work supports the degeneration of the PADI_N domain in cyanobacteria and demonstrates the existence of catalytically active PADI proteins in cyanobacteria [31]. The relatively recent appearance of the NTD during the PADI family evolution suggests it has a functional relevance that is unique to higher organisms.

Previously, we found that PADI2 interacts with P-TEFb to maintain the expression of actively transcribed genes [30,44]. The P-TEFb complex comprises cyclin-dependent kinase 9 (CDK9) and its regulatory partner cyclin T1 (CCNT1) [45]. Activation of the P-TEFb complex is required early in transcription to overcome RNAPII pausing and promote the productive phase of transcription elongation [45–48]. Lack of recruitment of the P-TEFb complex and associated components has been linked to several disease conditions [49–51], highlighting the importance of this process in healthy physiology. While PADI2 could be a therapeutic target that specifically prevents p-TEFb-mediated expression of genes that promote disease states, more needs to be understood.

Here, we analyze the recent evolution of the PADI protein family. We performed a comparative genomics analysis of PADI2 and identified 20 putative amino acid substitutions, in different species, that might be important for PADI2 structure and functionality in mammals. The majority of the selected amino acids are exposed in the three-dimensional (3D) structure and belong to the non-catalytic NTD or to the middle domain of PADI2. These positions suggest that they can participate in protein-protein interactions. We investigated the functional relevance of positively selected exposed amino acids in the NTD and middle domain of PADI2 by analyzing their impact on PADI2 interactions with the P-TEFb complex. We found that leucine 162 (L162), in the exposed loop of the middle domain of PADI2, underwent positive selection during the initial divergence of primates. Indeed, this residue contributes to interactions between PADI2 and the P-TEFb complex. These results establish a link between the evolutionary selection of key residues in PADI2 and their role in regulating the protein-protein interactions required for functional transcriptional regulation.

## Results

### Evolutionary analysis of the PADI family

To obtain a global picture of evolutionary relations within the PADI family, we first searched for all PADI family members with sequence homology to human PADI2 in the OMA (Orthologous MAtrix) database [52] and collected all the available sequences in mammals (see Materials and Methods). After close inspection, we discarded partial sequences and low-quality genomes, leaving 185 PADI orthologs that represent 25 mammalian species and three bird species (**S1 Fig**). We used these sequences to build a multiple sequence alignment (MSA, **S1 File**) and reconstructed a phylogenetic tree using the software RaxML, rooting the tree using the bird sequences as an outgroup (**S2 Fig,** see Materials and Methods).

The tree we built is consistent with five PADI genes that resulted from duplication of a common ancestral sequence. Indeed, each of the five well-defined clusters in the reconstructed tree corresponds to the five different *PADI* genes (*PADI1*, green; *PADI2* purple; PADI*3*, blue; *PADI4*, red; and *PADI6*, orange) with known species relationships being broadly recapitulated within each of these clusters (**S2 Fig)**. Robust bootstrap supports (82-100, **S3 Fig**) for this tree strongly support the hypothesis that the genes within each cluster are orthologous. We observed clustering between *PADI1* and *PADI3* and between *PADI4* and *PADI6*, with *PADI2* being the most distant ortholog among family members. Notably, PADI2 also has shorter branch lengths in comparison to other family members, indicating a higher level of conservation among family members in this cluster. PADI6 family members have the longest branch lengths, indicating this cluster is the most divergent and suggesting these genes arose from an old duplication event. Interestingly, there are only three PADI genes in each bird species, and only one set of clusters within the PADI2 family, suggesting that one or more duplication events took place after the divergence from birds. PADI2 is also the closest subfamily to the remaining PADI outgroup species clusters, indicative of an ancestral position in this family.

Two bird genes (i.e., ANAPL06996 and ANAPL06995 in duck) in the unpainted cluster (**S2 Fig**) did not exhibit clear orthology to specific *PADI* genes in either Ensembl or the OMA, though some were annotated as *PADI1* orthologs. Our data suggest that there was a bird-specific duplication, possibly from *PADI1,* which is the closest gene in the synteny analysis (**S4A Fig**). Our species comparison also revealed an inversion of the entire *PADI* locus that appears to have occurred in the common ancestor of humans and gibbons (Hominoidea; **S4B Fig**). Most mammalian species have the same gene order as that found in mice (**S4C Fig**). Overall, our analyses revealed key evolutionary changes at the *PADI* locus, including gene re-arrangements and duplications within mammals.

### PADI2 is highly conserved across species

Since PADI2 is likely to be the ancestral copy, we decided to focus on the most recent changes occurring in its sequence. By focusing on a specific phylogenetic selection of species, we aimed to identify changes that have occurred since the common amniote ancestor that occur with good resolution in different mammals. Our dataset contains *PADI2* representatives in all three major extant groups in class Mammalia and the principal orders or infraclasses: primates, rodents, carnivores, cetartiodactyla, lagomorphs, proboscidea, and marsupials. We built a multiple sequence alignment with all 28 *PADI2* sequences (see Materials and Methods**, S1 File)** and computed the ratio of substitution rates at non-synonymous sites and synonymous sites (dN/dS; also termed Ka/Ks) to infer the direction and magnitude of natural selection acting on protein-coding genes [53]. The difficulty of estimating the dN/dS ratio grows with the phylogenetic distance between sequences. Maximum likelihood methods were used to correct for multiple substitutions in the same site, and we estimated important parameters, such as the divergence between sequences or the transition/transversion ratio by deducing the most likely values to produce the input data [53].

We calculated the dN/dS ratio for all internal branches and leaves in the tree using CodeML (**Fig 1**). We observed low dN/dS values whereby most of the branches had a value of < 0.2. This was consistent with negative selection; in other words, changes in this genetic sequence were actively selected against. This analysis supports the idea that PADI2 is a highly conserved protein, suggesting that it performs critical functions in the organism (**Fig 1**). However, we observed a relatively high dN/dS in the common ancestor of all primates, possibly indicating relaxation of selection or the fixation of a few beneficial mutations due to positive selection at this point in evolution. The dN/dS ratio requires a high number of changes to detect a selection effect and our data indicate that dN/dS values are very low in PADI2. Another limitation of this method is that it does not distinguish amino acid substitutions that are chemically similar from others with very different properties. However, this analysis suggests that intriguing selection processes operated during PADI2 evolution.

**Fig 1 (related to S1-S4 figs and File S1).**
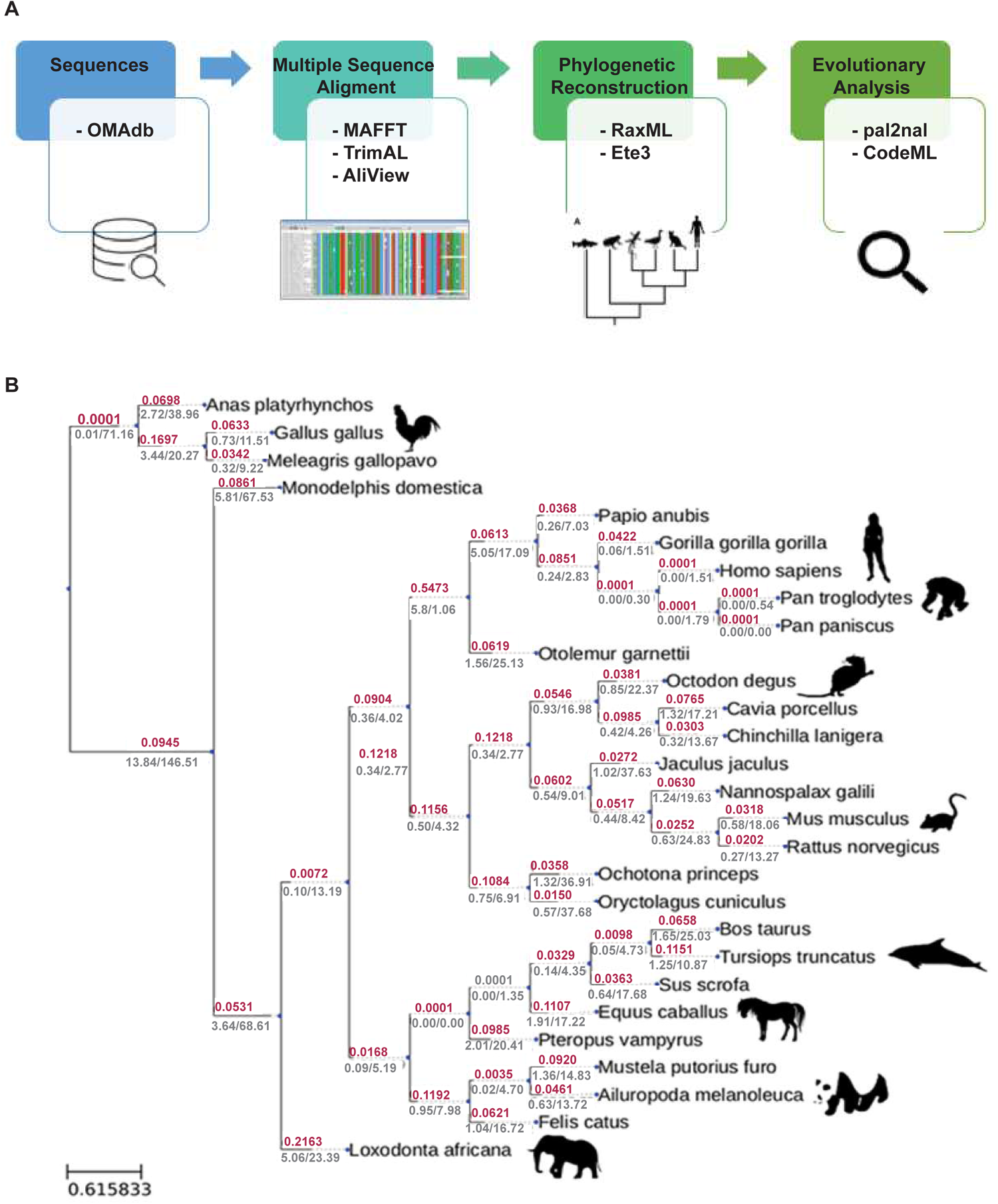
PADI2 conservation across species. **(A)** Diagram showing the main steps involved in phylogenetic reconstruction and evolutionary analysis. Each box contains a list of the tools used. (**B**) The PADI2 gene tree showing the dN/dS ratios (red) for every branch as well as dN and dS (grey) for every branch. The common branch to all primates is highlighted in blue. Branch lengths are proportional to nucleotide substitutions, as calculated by ete3.

### Detection of positively selected residues in PADI2

We used an orthogonal approach, the branch site positive selection test, to surpass the limitations of dN/dS analysis and identify specific residues that have undergone selection [54,55]. This test can discriminate between the relaxation of selection or positive selection and can detect individual residues that are under positive selection in a particular lineage. This test compares a null model in which codon-based dN/dS for all branches can only be ≤ 1; in the alternative model, the labeled foreground branch may include codons evolving at dN/dS > 1 [54]. This test can discriminate between the relaxation of selection or positive selection and can detect individual residues that are under positive selection in a particular lineage. After running the branch-site positive selection test on every branch of the PADI2 tree, we detected 20 individual relevant substitutions in different parts of the tree. We removed a few candidates for positively selected amino acids detected by CodeML in *Loxodonta africana* due to the alignment of a non-homologous region (238-278), possibly due to an assembly error in that species. The residues identified in the different species or branches are shown in **Table 1 and Fig 2**.

**Fig 2 (related to Table 1).**
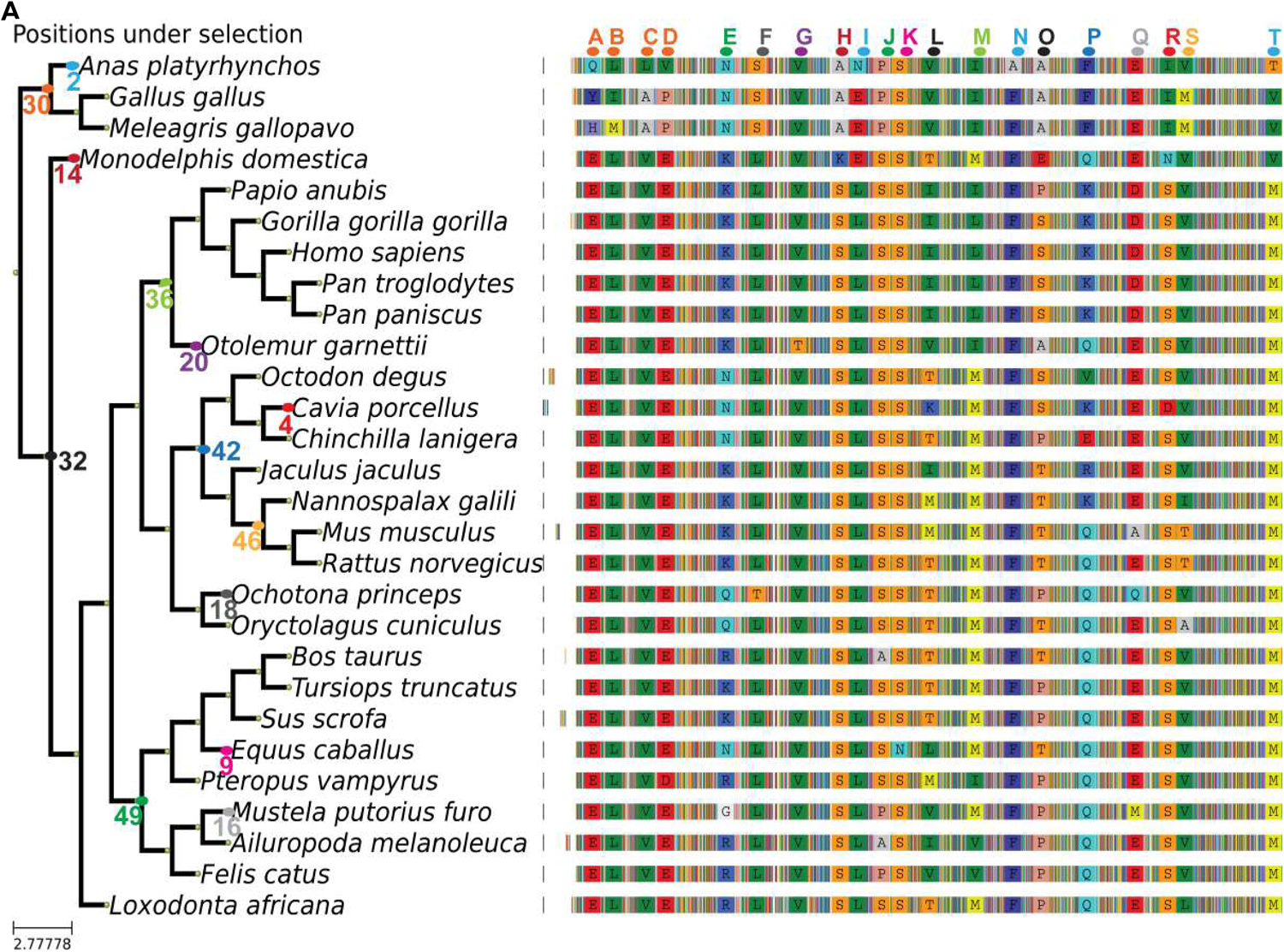
Summary of the branch site test results for internal and terminal branches. (**A**) Summary of the multiple sequence alignment, showing only columns with positions that are detected as being under positive selection with the branch site test. The colored-blurred positions in between represent the rest of the alignment (illustrated in Supplementary File 1). The node number correspondence can be found in Supplementary Figure S1.

**Table 1.** **Positively** selected sites across species under the branch site (CodeML) and mapped onto the PADI2 human structure. The position of the amino acid in the domain, MSA, and PDB file (Supplementary material file S2) are shown in the domain, sequence position, and PDB number, respectively. The flanking residues are shown with the given positively selected amino acid depicted in bold. The type of secondary structure and solvent accessibility (i.e., exposure) is shown in the secondary structure and accessibility column. Finally, the MVORFFIP scores (see Materials and Methods) are also indicated.

| Domain | Alignment position | Sequence position (human) | Secondary structure | Flanking residues | MVORFFIP scores (0-1) | Accessibility |
| --- | --- | --- | --- | --- | --- | --- |
| PADI_N | 70 | E16 | Beta strand | RVEAV | 0.8 | Exposed |
| PADI_N | 79 | L25 | Beta strand | TYLWT | 0.0 | Buried |
| PADI_N | 107 | V53 | Beta strand | EVVRD | 0.5 | Exposed |
| PADI_N | 114 | E60 | Loop | AEEVA | 0.7 | Exposed |
| PADI_N | 190 | K136 | Loop | NP**K**KA | 0.6 | Exposed |
| PADI_N | 216 | L162 | Loop | PWL**P**K | 0.7 | Exposed |
| PADI_N | 260 | V201 | Beta strand | EIVLY | 0.2 | Exposed |
| PADI_N | 304 | S245 | Loop | GG**S**AE | 0.6 | Exposed |
| PADI_N | 308 | L249 | Beta strand | ELLFF | 0.8 | Exposed |
| PADI_N | 322 | S263 | Loop | GF**S**GL | 0.7 | Exposed |
| PADI_N | 326 | S267 | Beta strand | LV**S**IH | 0.7 | Exposed |
| PADI_N | 350 | T289 | Beta strand | TD**T**VI | 0.6 | Exposed |
| PADI_C | 401 | L342 | Loop | QY**L**NR | 0.7 | Partially buried |
| PADI_C | 439 | F380 | Loop | KD**F**PV | 0.8 | Exposed |
| PADI_C | 460 | S401 | Loop | FE**S**VT | 0.8 | Exposed |
| PADI_C | 511 | K452 | Helix | FL**K**AQ | 0.7 | Exposed |
| PADI_C | 567 | D507 | Helix capping | QK**D**GH | 0.5 | Exposed |
| PADI_C | 596 | S536 | Helix | NE**S**LV | 0.7 | Exposed |
| PADI_C | 598 | V538 | Helix | SL**V**QE | 0.8 | Partially buried |
| PADI_C | 723 | M663 | Helix capping | WH**M**VP | 0.7 | Partially buried |

### Positively selected residues in PADI2 may mediate protein-protein interactions

We next integrated our evolutionary insights into PADI2 with known structural information to study the properties of positively selected residues. To pursue this analysis, we mapped all the positively selected amino acids onto the structure of PADI2. 12 of the 20 positively selected amino acids mapped onto the PADI2_N and PADI2_M domains, while eight mapped to the PADI2_C domain. The latter contains the catalytic pocket and is responsible for the citrullination of arginine residues. (**Table 1**, **Fig 3A-B**). Of note, only one of the positively selected amino acids (L25) is fully buried (amino acids are numbered as shown in **Table 1, File S2, Movie S1**); most of the other amino acids are fully exposed except for L342, V538, and M663, which are partially buried. In the secondary structure, approximately 50% of the amino acids are located in loops, while the others are either in beta strands or alpha helices (including helix-capping regions) (**Table 1**).

**Fig 3 (related to Table 1, File S2 and Movie S1).**
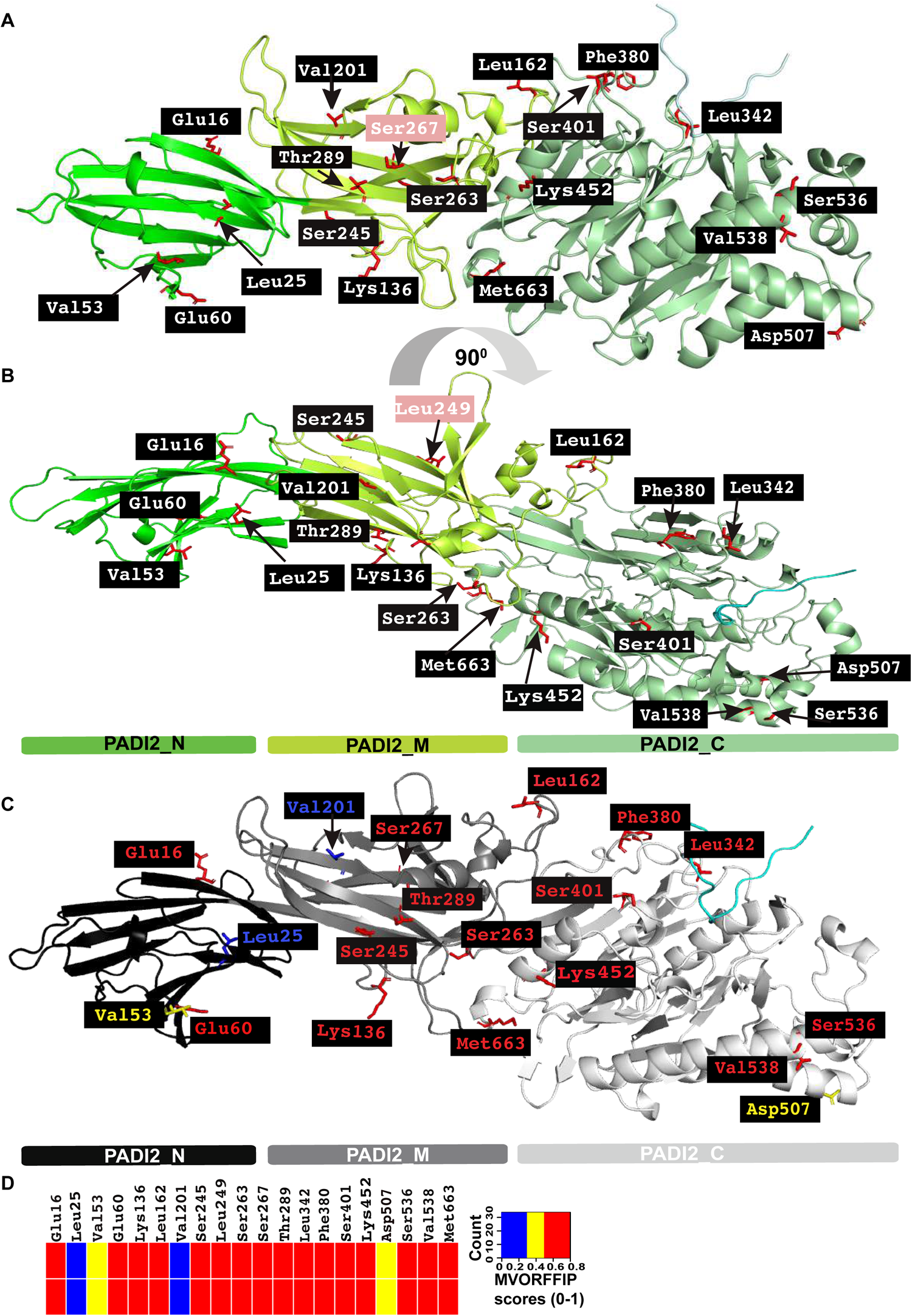
Mapping of positively selected amino acids onto the PADI2 structure. **A-B.** Ribbon representation of PADI2 showing the N-terminal domain (PADI2_N), the middle domain (PADI2_M), and the C-terminal domain (PADI2_C) in different shades of green (in green, bright green, and pale green, respectively). (**B)** The same as (A) but applying a 90-degree rotation over the longitudinal axis. Pink highlighted residues are specific to the respective orientation. PyMol session of PADI2 and the highlighted amino acids are available in Supplementary Material File S2. (**C**) Cartoon representation of PADI2 with positively selected amino acids colored according to MVORFFIP scores as shown in the heatmap in panel D; PADI2_N, PADI2_M and PADI2_C domains colored in black, grey, and white, respectively. (**D**) The heatmap is based on the MVORFFIP scores as mentioned in Table 1.

After evaluating the features of PADI2_N and PADI2_M domains and analyzing each particular amino acid and its potential role based on structural data, we concluded that the likely roles of E16, V53, E60, K136, L162, V201, S245, S249, S267, and T289 are to mediate interactions with putative binding partners. These amino acids are fully exposed, and their chemical properties correspond with classical protein interfaces [56,57]. We thus characterized these positively selected amino acids using Multi-VORFFIP (MVORFFIP), a tool that predicts protein-, peptide-, DNA-, and RNA-binding sites [56]. Notably, the MVORFFIP predictions for all except V201 and L25 yielded very high scores (> 0.7, on a scale of 0 to 1), supporting the hypothesis that they are interface residues (**Fig 3C-D**, **Table 1**). S263 is located in a long loop close to the interface between domains, suggesting that it plays a role in hinge motions and geometrical orientation between domains. Given its position in the structure, the likely role of L25 (the only positively selected amino acid that is completely buried) may be to prevent distortion of a nearby beta-sandwich. Indeed, a larger hydrophobic amino acid at this position would change the packing of the beta-strands.

We found that three positively selected residues in the PADI2_C domain, L342, F380, and S401, are close to the catalytic pocket. These residues are likely to play a role in substrate specificity (L342) and in the dynamics of catalysis (F380 and S401). K452, S536, and V538, however, are located in a helix that is distant from the active site pocket. Judging from the structural microenvironment, these residues could also play a role in protein-protein interactions, in line with the high scores assigned by MVORFFIP (**Fig 3D**, **Table 1)** [56]. Finally, D507 and M663 are both located in helix-capping positions. Conservation in helix capping is important for the stability and integrity of the helix. Residue M663 may also play a role in the packing of the C-terminal tail. A CAPS Database search [58] showed that other helices with the same capping structure present a conserved small hydrophobic patch in the same position by including M, V, or L residues. It is noteworthy that M663 is indeed strictly conserved across all human PADIs. These observations suggest that most of these positively selected amino acids may function in facilitating and stabilizing protein-protein interactions.

### The middle domain of PADI2 maintains interactions with the P-TEFb complex

We next tested whether residues that have been positively selected in evolution but are not linked to the catalytic domain of PADI2, might interact with other proteins. For this analysis, we examined the P-TEFb kinase complex, which we previously found to interact with PADI2 [30]. The structures of PADI2 [25] and CDK9-CCNT1 [59] are known individually, and therefore it was possible to derive a structural model of the PADI2–CDK9/CCNT1 complex by protein docking. After deriving the docking ensemble of PADI2-CDK9/CCNT, we ranked the models. Importantly, we used a swarm-based approach (among other metrics) to select the putative model of interaction, reasoning that determining the region of PADI2 would be relevant to its interaction with CDK9/CCNT1 (**S4D Fig, File S3, Movie S2**). Moreover, we derived the structural model using AlphaFold-Multimer [60,61]. As shown in **S4E Fig (File S4, Movie S3)**, interacting regions in the structural model from AlphaFold-Multimer overlap substantially with those identified by docking (**S4D Fig**). More specifically, the predicted interface between PADI2 and CDK9-CCNT1 contained PADI2_M and the hinge region was overrepresented with the number of positively selected residues. Importantly, L162 is present on the highly exposed loop (**Fig 4A**).

**Fig 4 (related to File S5, Movie S4, S4 Fig).**
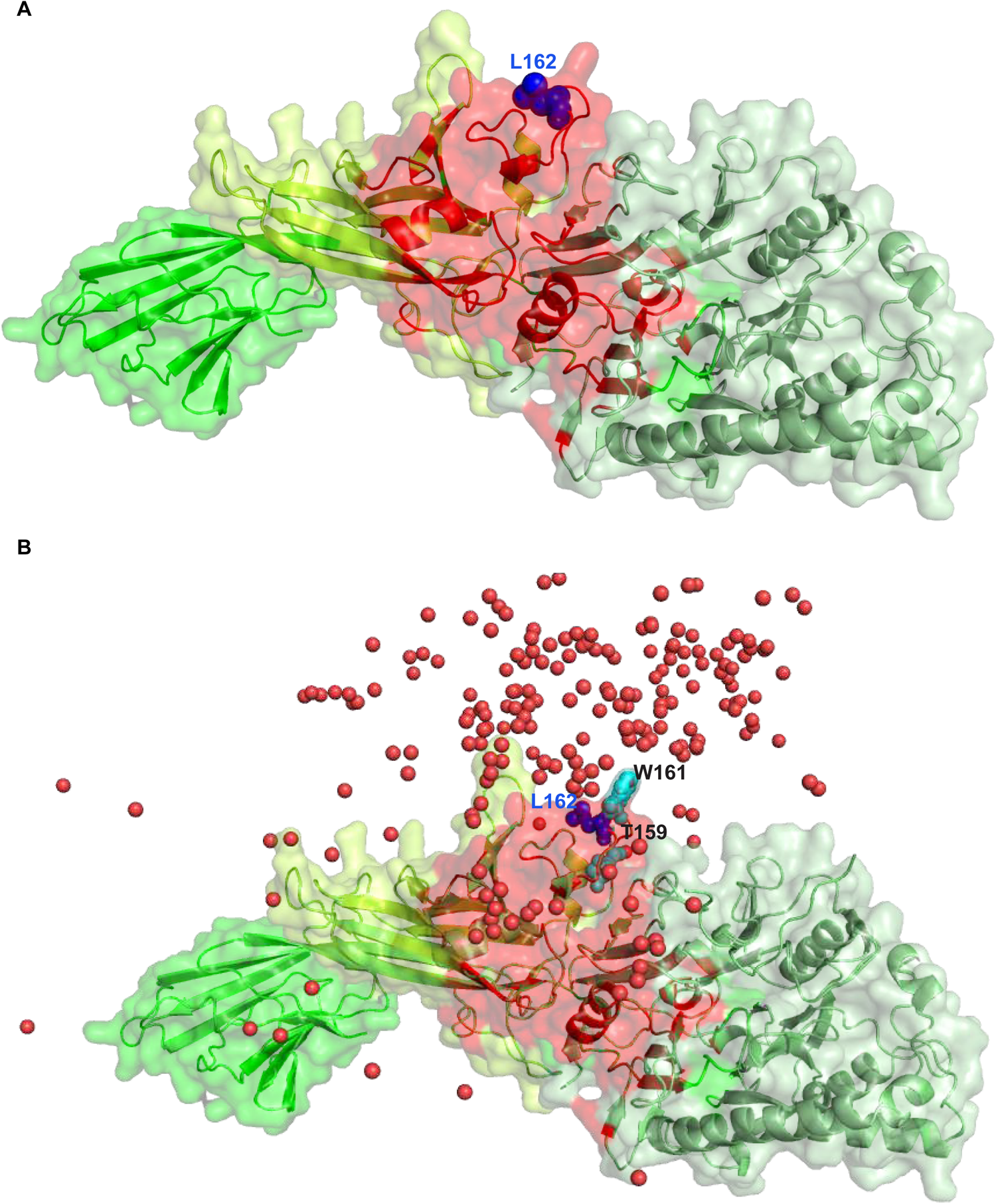
The predicted interface between PADI2 and the P-TEFb complex. **(A)** Surface representation of PADI2 with the same color scheme as in Figure 3 with the consensus interface with P-TEFb complex derived from docking and AlphaFold-Multimer structural models highlighted in red and L162 shown as a blue sphere. (**B**) The same representation as in (A) including the top 200 docking poses represented as red spheres depicting the center of mass of CDK9/CCNT1; T159 and W161 are also shown in cyan as sphere representation. The PyMol session of PADI2 with the P-TEFb complex with docking and AlphaFold-Multimer shared region is available in Supplementary Material File S5.

While AlphaFold-Multimer generates a single model, docking is represented by a range of docking poses. This range of models allowed us to identify regions of the predicted interface, i.e., PADI2_M and PADI2_C (**Fig 4A**), that are over-represented in the docking space. The distribution of the top 200 docking poses (represented using a unique point depicting the center of mass of CDK9/CCNT1) revealed that the region around the hinge between the PADI2_M and PADI2_C was over-represented **(Fig 4B, File S5, Movie S4)**. This particular region includes L162, a residue overrepresented in the docking conformers, along with T159 and W161. These three residues are located in a highly exposed region. We therefore selected these residues for experimental validation. Note that L162 in PADI2 was present among the top-ranking interface residues likely to meditate the interaction with CDK9/CCNT1 and is also a positively selected residue, suggesting that it is important for maintaining this interaction.

### The loop encompassing positively selected Leu162 is important for cell proliferation and maintaining PADI2 interactions with the P-TEFb complex

Since the P-TEFb kinase complex is involved in transcription elongation and cell proliferation [62–64], we experimentally tested whether the L162-encompassing loop in the middle domain of PADI2 affects cell proliferation. In addition to L162, we tested its neighboring amino acids, W161 and T159, which also ranked highly in our evolutionary and interaction prediction analysis and are located in a highly exposed loop region structurally close to the boundary between the PADI2_M and the catalytic domain. We selectively expressed the green fluorescence protein (GFP)-tagged wild-type (WT) PADI2 as well as the single (T159A or W161A or L162A), double (L162A/W161A), or triple (L162A/W161A/T159A) mutant of PADI2. The GFP-positive cells were sorted by FACS to ensure positive cell selection (**S5A-B Fig**). We monitored cell proliferation in HeLa cells expressing the GFP-tagged PADI2 WT and mutants (single, T159A or W161A or L162A, double, L162A/W161A, and triple, L162A/W161A/T159A). Strikingly, we observed that among the three single PADI2 mutants, L162A significantly decreased cell proliferation compared to WT, highlighting the functional relevance of L162 in the middle domain of PADI2 (**Fig. 5A, S5C Fig).** In the same manner, the PADI2 double mutant L162A/W161A, as well as the triple mutant L162A/W161A/T159A, showed a further significant reduction in cell proliferation. These observations highlight the functional role of L162A in this remarkably exposed loop in the middle domain of PADI2.

**Fig 5 (related to S5).**
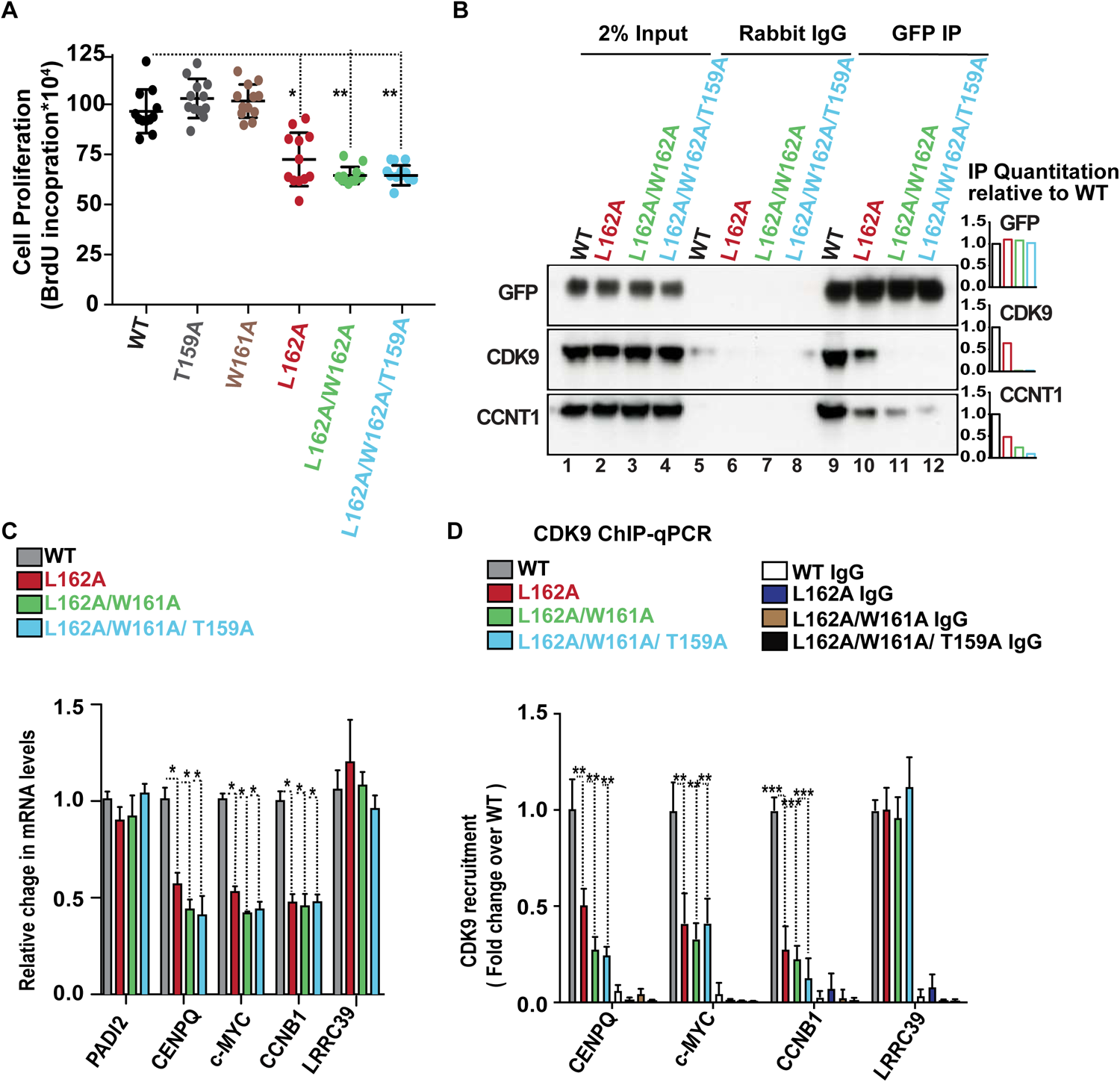
Positively selected L162 encompassing loop contributes to its interaction with the P-TEFb complex. (**A**) Cell proliferation of HeLa cells specifically expressing the WT or mutant PAD12 (single, T159A or W161A or L162A; double, L162A/W161A or triple, L162A/W161A/T159A). Data represent mean ± SEM of at least eight biological experiments. *p value < 0.05; **p value < 0.01. **(B**) Immunoprecipitation with GFP-specific antibody or non-immune rabbit IgG of GFP positive Hela cells nuclear extracts expressing wild-type (WT)or mutant (single, L162A; double, L162A/W161A or triple, L162A/W161A/T159A) PADI2 followed by western blot with the indicated antibodies. The relative quantification is shown as a bar plot. (**C**). Quantitative RT-qPCR validation in HeLa cells selectively expressing the WT or mutant PAD12 (as in B). Changes in mRNA levels were normalized to *GAPDH* mRNA. Data represent the mean ± SEM of n ≥ 3 biological experiments for all plots in the figure. Two-tailed unpaired Student’s t-test was used to determine the statistical significance between the groups. (**D**) ChIP-qPCR assay performed in HeLa cells selectively expressing the WT or mutant PAD12 (as in B) with CDK9 antibody. Non-immune IgG was used as a negative control. Y-axis: fold change over the input samples. Data represent mean ± SEM of three biological experiments, *p-value < 0.05; **p value < 0.01.

Therefore, we focused on the L162A, L162A/W161A, and L162A/W161A/T159A mutants along with WT PADI2 for further analysis. In the immunoprecipitation assay, using a GFP-tagged antibody, we observed that the P-TEFb complex (containing CCNT1 and CDK9) from HeLa cells was efficiently immunoprecipitated with GFP-WT PADI2 but not with GFP-L162A-PADI2 (**Fig 5B).** These results confirmed the functional role of the positively selected L162 in maintaining PADI2 interactions with the P-TEFb complex. Likewise, the P-TEFb complex was only weakly immunoprecipitated with either the double or triple PAD12 mutant, highlighting the function of the highly exposed region in the middle domain of PADI2 (PADI2_M) in this protein-protein interaction. Considering that proper recruitment of the P-TEFb complex is required for efficient transcription of highly expressed genes relevant to cell proliferation, we validated the reduced expression levels of the *c-MYC*, *CENPQ*, and *CCNB1* genes in cells expressing the single, double, or triple PADI2 mutant (**Fig 5C)**. Of note, the levels of PADI2 expression did not differ significantly. We observed that the L162A single mutant (as well as double and triple mutants) significantly and specifically reduced the expression of highly expressed genes, including *c-MYC*, and *CCNB1, CENPQ* without affecting the levels of a control gene (the low-expressed *LRRC39* gene). Next, we examined if the L162A mutant in PADI2 can mediate CDK9 recruitment to the promoter region of the target genes. We performed a chromatin immunoprecipitation (ChIP) assay using a CDK9-specific antibody in HeLa cells specifically expressing either WT PADI2, the L162A mutant, or the double and triple mutants. We found that CDK9 occupancy decreases in the presence of mutants compared to the WT PADI2 control (**Fig 5D)**, suggesting that the positively selected L162 residue, along with W161A/T159A, is important to maintaining PADI2 interaction with the P-TEFb complex.

Given the fact that the L162-encompassing loop is important to maintaining PADI2 interaction with the P-TEFb complex, we next explored the effects of the PADI2 triple-mutant (L162A/W161A/T159A) on the transcriptome using single-cell RNA sequencing experiments. We compared the single-cell transcriptomics atlas of PADI2 triple-mutant to that of WT PADI2-expressing HeLa cells (**S6A-B Fig).** After quality control, 49,503 cells were subjected to downstream analysis, of which, 17,139 and 32,364 cells were derived from the WT and triple mutant PADI2 samples, respectively. We did not observe any significant differences in cell populations through unsupervised clustering and dimensionality reduction. TSNE plots organized the data into three clusters (**S6A Fig).** However, these clusters are not significantly different in triple mutants in comparison to WT (**S6B Fig).** Among the differentially expressed genes, we found *c-MYC* (**Fig 6A-B)**. Within the top genes that are differentially expressed, we also found *CCNB1* and *CENPQ,* which are related to cell cycle regulation (**Fig 6A-B)**. Remarkably, in global differential expression (DEseq) analysis, the PADI2 triple mutant (L162A/W161A/T159A) affected the expression of over 469 genes (Log2FC>0.58 or Log2FC<-0.58, p-value <0.05). Of these, 162 genes were downregulated and 307 upregulated (**Table S2).** Gene ontology analysis of the top down-regulated genes (Log2FC<-1.0, p-value <0.01) revealed enrichment in the cell cycle, RNA binding, and chromosome organization. The up-regulated genes were instead enriched for genes linked to the inflammatory response, cell-cell signaling, and cell adhesion processes (Log2FC>1.0, p-value <0.01) (**Fig 6C and Table S3)**. We also investigated the regulatory gene sets using the gene ontology tool. The promoter regions of downregulated genes were significantly enriched for GTF2A2 (UniProt: P52657), GTF2E2 (UniProt: P29084), FOXE1 (UniProt: O00358), MZF1(UniProt: P28698), ZFHX3_(UniProt: Q15911) binding motifs (**Fig 6D and Table S4**). Meanwhile, the upregulated genes showed significant enrichment in the promoter regions for motifs that can bind ZNF92 (UniProt: Q03936), METTL14 (UniProt: Q9HCE5), GFI1 and CTTTGA motif (**Fig 6E and Table S4**). This analysis highlights a connection between regulatory elements and differentially expressed (WT vs PADI2 triple mutant) genes.

**Fig 6 (related to S6).**
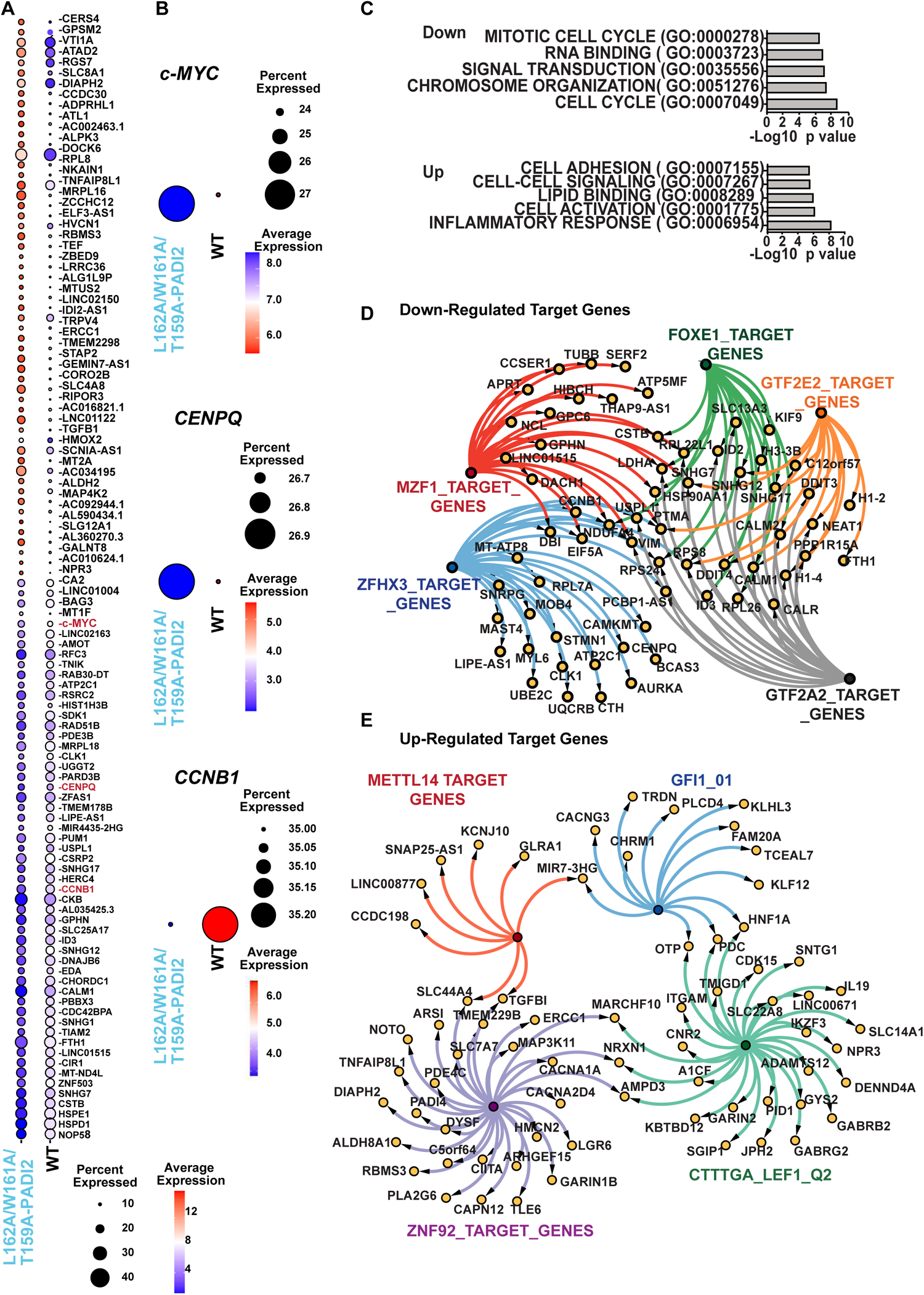
The L162 enclosed loop regulates the transcription regulation. (**A**). Dotplot depicting the top 55 up- and downregulated genes in L162A/W161A/T159A-PADI2 mutants vs. WT. Selected were the p_adj_ <0.05 and highest FC (fold change) according to the Find Markers function in Seurat (4.3.0, log2FC.threshold=0.25, min. pct=0.1). (**B**). Dot plots depicting the expression of c-MYC and CCNB1 in L162A/W161A/T159A-PADI2 mutants vs. WT. The circle size depicts the percentage of cells expressing the respective gene, while the color encodes the average expression. (**C**). Gene set enrichment analysis (GSEA) for molecular functions and biological processes. Representative processes are presented (*Upper* -Downregulated genes, *Lower*-Up-regulated genes). The X-axis shows the -log10 transformed p-values. GO, Gene Ontology. (**D-E**). The connection map of the target genes from the representative regulatory elements in WT/ (L162A/W161A/T159A) PADI2 (**D**) upregulated genes (**E**) downregulated genes.

Importantly, *c-MYC, CCNB1,* and *CENPQ* overexpression have been linked to oncogenesis [65–70]. These results support the functional relevance of the positively selected L162-encompassing loop in the middle domain of PADI2 in the expression of important oncogenic genes. Modulation of *c-MYC* expression by epigenetic regulation in cancer cells has been suggested to increase heterogeneity in the transcriptional landscape that promotes tumorigenesis [71]. These observations suggest a potential function for the L162-encompassing loop of PADI2 in the regulation of the *c-MYC /CCNB1/CENPQ* axis which has been linked to tumorigenesis.

## Discussion

We characterized the evolution of the PADI gene family using the most complete dataset of mammalian orthologous sequences to date. The three major extant groups in class Mammalia are organized into two subclasses: Prototheria which includes monotremes (platypus and echidnas), and Theria, which includes the infraclasses Methatheria (marsupials), and Eutheria (placental mammals). According to Jones et al. 2009 [72], there are 5416 mammalian species divided into 29 orders. Most known species belong to the 7 biggest orders (4865 species), all of which have at least one representative in our phylogenetic analysis.

The functional relevance of PADI2 is highlighted by its high sequence conservation across species and very low dN/dS values. We took advantage of the natural experiments performed by evolution and applied sensitive methodologies such as the branch-site test of positive selection. Comparing the amino acid sequences of 25 mammalian species, we identified 20 positively selected residues predominantly located in the non-catalytic domain of the PADI2 in different species and phylogenetic branches **(Figs 1-2, S1-S4 Figs)**. Since these selected residues could have a strong functional impact on PADI2, we studied them further using the human sequence.

By examining the location of the positively selected residues in the structure of human PADI2, modeled in a complex with R1810 in the CTD of RNAPII, we determined that the majority of positively selected PADI2 residues were structurally highly exposed in the non-catalytic domains of PADI2 (**Fig 3**, **Table 1**). Notably, the highly exposed position and chemical properties of these positively selected residues suggested they play important functions in cellular processes. In addition, our MVORFFIP analysis revealed that these interface residues may play a significant role in protein-protein interactions (**Fig 3D**). Therefore, it could be postulated that the appearance of these positively selected residues contributed to the evolution of PADI2 function by modulating its interaction with an essential cognate set of proteins. Further work will be required to investigate these possibilities.

Specifically, the P-TEFb complex, comprising CCNT1 and CDK9, interacts with PADI2 and facilitates the effects of citrullination of RNAPII-R1810 on transcription [30]. Hence, we used structure modeling approaches to derive the tertiary structure of the PADI2–CCNT1/CDK9 complex. To identify the PADI2 residues that contribute to forming the PADI2–CCNT1/CDK9 complex, we analyzed structural models and identified a highly exposed loop in the PADI2_M domain as a potential interacting region. Strikingly, this loop includes the positively selected residue L162, along with T159 and W161. This observation supports the notion that positively evolved residues tend to coincide with the important residues on the protein surface, and that positively selected structural clusters are important for cellular function (**S6C Fig**).

Our data in HeLa cells overexpressing the GFP-tagged WT and a mutant PADI2 also unveiled an essential function for the L162-containing loop in interaction with the P-TEFb kinase complex and cell proliferation. Notably, mutation of L162, alone or in combination with mutated W161 and/or T159, reduced (i) cell proliferation, (ii) PADI2 interaction with P-TEFb kinase complex, and (iii) reduced the PADI2-dependent expression of highly expressed genes that are relevant for cell growth. Note that one limitation of our study was that we could not mutate the endogenous PADI2 gene due to the hyperpolyploid nature of the HeLa genome. Therefore, we based our analysis on overexpression of GFP-tagged wild-type and mutant versions of PADI2 in HeLa cells.

We focused on the interactions between PADI2 and the P-TEFb complex, specifically with the positively selected L162-encompassing loop. However, in the future, it would be interesting to investigate the functional role of other positively selected PADI2 amino acids in cellular processes. Nonetheless, our analysis shows the importance of using a multi-disciplinary approach. Here, we used comparative genomics, evolutionary analysis, structural modeling, and genetic perturbations to analyze the functional relevance of a recently evolved non-catalytic domain in a highly conserved enzyme family. Overall, this evolutionary approach led to the identification of a PADI2 structural loop that is under positive selection and specifically supports an important role for PADI2 in the modulation of transcription elongation.

## Materials and Methods

### Cell lines

HeLa cells (ATCC CCL-2) were grown in DMEM with 10% FBS and 100U/ml penicillin-streptomycin according to the ATCC’s recommendations. Cells were transfected using Lipofectamine 3000 (Invitrogen) according to the manufacturer’s instructions.

### PADI2-GFP plasmids and fluorescence-activated cell sorting (FACS) sorting

PADI2 was cloned into the pCPR0032 GFP-tagged vector using the forward primer (F) 5′-AGAACCTGTACTTCCAATCCATGCTGCGCGAGCGGAC-3′ and the reverse primer (R) 5′-GATCCGTATCCACCTTTACTTTAGGGCACCATGTGCCACC-3′ using Gibson assembly [73]. The selected plasmid sequence was verified by Sanger sequencing. Next, PADI2 wild-type (WT) was used to generate a single mutant (T159A or W161A or L162A), a double mutant (L162A/W161A), and a triple mutant (L162A/ W161A/ T159A), using the following primers:

### PADI2-T159A

F1 5′-TGGTGAACTGTGACCGAG**AGG**CACCCTGGTTGCCCAAGGAGGACTGCCGTGATG −3′ R1 5′-CATCACGGCAGTCCTCCTTGGGCAACCAGGGTG**CCT**CTCGGTCACAGTTCACCA −3′.

### PADI2-W161A

F1 5′-TGGTGAACTGTGACCGAGAGACACC**CGC**GTTGCCCAAGGAGGACTGCCGTGATG −3′ R1 5′-CATCACGGCAGTCCTCCTTGGGCAAC**GCG**GGTGTCTCTCGGTCACAGTTCACCA −3′.

### PADI2-L162A

F1 5′-TGGTGAACTGTGACCGAGAGACACCCTGG**GCA**CCCAAGGAGGACTGCCGTGATG-3′ R1 5′-CATCACGGCAGTCCTCCTTGGG**TGC**CCAGGGTGTCTCTCGGTCACAGTTCACCA-3′.

### PADI2-L162A/ W161A

F2 5′-TGGTGAACTGTGACCGAGAGACACCC***GCC*GCA**CCCAAGGAGGACTGCCGTGATG −3′ R2 5′-CATCACGGCAGTCCTCCTTGGG**TGC*GGC***GGGTGTCTCTCGGTCACAGTTCACCA −3′.

### PADI2-L162A/W161A/T159A

F3 5′-TGGTGAACTGTGACCGAGAG**<u>GCA</u>**CCC***GCC*GCA**CCCAAGGAGGACTGCCGTGATG −3′ R3 5′-CATCACGGCAGTCCTCCTTGGG**TGC*GGC***GGG**<u>TGC</u>**CTCTCGGTCACAGTTCACCA-3′.

The corresponding fragments were generated by each pair of primers using oligonucleotide assembly and introduced by Gibson assembly [73]. All the generated mutants were confirmed by Sanger sequencing.

For transfection, 2 × 10^6^ HeLa cells were seeded in 10-cm plates, and 4 ug of the plasmid being tested was transfected using Lipofectamine 3000 (Invitrogen) for 24 hours according to the manufactureŕs instructions. Cells were trypsinized, and GFP-positive live cells were sorted using BD influx (Becton and Dickinson, San Jose, CA). Briefly, cells were stained with 1μg/mL concentration of DAPI (4′, 6-diamidino-2-phenylindole) before FACS sorting. A SSC-H (side scatter height) versus FSC-H (forward scatter height), morphological-related parameters dot-plot was used to exclude debris by gating cells; doublets were then excluded using a FSC-H versus FSC-A (forward scatter area) by gating and dead cells were excluded using DAPI versus FSC-A dot-plot by gating living cells. GFP-positive cells were identified and isolated using a G4 gate in GFP versus autofluorescence (AF) dot-plot. Obtained data were analyzed using Flow Jo 10. 6. GFP-positive cells were used for experiments. Cells were centrifuged and stored as pellets at –80°C prior to RNA extraction and immunoprecipitation experiments.

### Cell Proliferation assay

#### BrdU (5′-bromo-2′-deoxyuridine) cell proliferation assay

HeLa cells (0.3 × 10^3^) were transfected with either wild-type PADI2 (WT) or mutant PADI2 (with L642A or T159 or W161A or L642A/W161A, or L642A/W161A/T159A) using Lipofectamine 3000 (Invitrogen) in 96-well plates. The cell proliferation ELISA BrdU (5′-bromo-2′-deoxyuridine) colorimetric assay (Roche,11647229001) was performed as per the manufacturer’s instructions. The experiments were performed on at least eight biological replicates.

### Incucyte ® Proliferation Assays for Live-Cell Analysis

HeLa cells seeded in 96-well plates with 300 cells per well, transfected with either wild-type PADI2 (WT) or mutant PADI2 (with L642A or T159 or W161A or L642A/W161A, or L642A/W161A/T159A). After 24hours of transfection, imaging was performed using the IncuCyte live cell imaging system (Essen BioScience). Scans at 4× magnification were taken every 8 hours for 5 days. Cell confluence was calculated from microscopy images using the Incucyte software algorithm to generate a proliferation index corresponding to the change in confluence for each well. These measurements are the mean ± SEM of at least six replicates.

### RNA extraction and RT-qPCRs

RNA from HeLa cells transfected with either wild-type PADI2 (WT) or mutant PADI2 (with L642A, or L642A/W161A, or L642A/W161A/T159A) was extracted using RNeasy (Qiagen) according to the manufactureŕs instructions. Purified RNA (1µg) was used for DNase treatment (Thermo Scientific) and quantified with a Qubit 3.0 Fluorometer (Life Technologies).

Reverse transcription of RNA was performed using a qScript™ cDNA Synthesis Kit (Quanta Bioscience 95047-100) according to the manufactureŕs instructions. Complementary DNA was quantified by qPCR using Roche Lightcycler (Roche), as previously described [74]. For each gene product, relative RNA abundance was calculated using the standard curve method and expressed as relative RNA abundance after normalizing against the human *GAPDH* gene level. All gene expression data generated by RT-qPCR represented the average and ± SEM of at least three biological replicates. Primers used for RT-qPCR are listed in **Table S1**.

### Single-cell samples, library preparation, and sequencing

HeLa cells (0.6 × 10^6^) were transfected with either wild-type PADI2 (WT) or the PADI2 triple mutant (with L642A/W161A/T159A) using Lipofectamine 3000 (Invitrogen) in 60mm plates. GFP-positive singlet cells and live cells were suspended in 1ml of PBS+BSA 0.05%. Cells were centrifuged at 400 rcf for 5 min at 4°C in order to bring the cell concentration to 300-1000 cells/µl. Cell concentration and viability were determined by manual counting using a Neubauer chamber and staining the cells with Trypan blue. Cells were partitioned into Gel Bead-In-Emulsions (GEMs) using the Chromium Controller system (10X Genomics), with a target recovery of 5000 total cells per sample. cDNA sequencing libraries were prepared using the Next GEM Single Cell 3’ Reagent Kits v3.1 (10X Genomics, PN-1000268), following the manufacturer’s instructions. Briefly, after GEM-RT clean-up, cDNA was amplified for 11 cycles, and cDNA QC and quantification were performed on an Agilent Bioanalyzer High Sensitivity chip (Agilent Technologies). cDNA libraries were indexed by PCR using the PN-1000215 Dual Index Kit TT Set A Plate. Size distribution and concentration of 3’ cDNA libraries were verified on an Agilent Bioanalyzer High Sensitivity chip (Agilent Technologies). Finally, sequencing of cDNA libraries was carried out using the Illumina NovaSeq 6000.

### Single-cell RNA sequencing data processing and analysis

Transcriptome-wide analysis of human L162A/W161A/T159A-PADI2 mutant and WT cells was performed at single-cell resolution using NovaSeq 6000 pipeline. Sequencing data were aligned to the human reference genome Grch38. Data with at least 500 unique molecular identifiers (UMIs), log10 genes per UMI >0.8, >250 genes per cell and a mitochondrial ratio of less than 20% were extracted, normalized, and integrated using the Seurat package v4.0.3 in R4.0.2. After quality control and integration, we performed a modularity-optimized Louvain clustering. After quality control, 17,139 WT and 32,364 PADI2mut cells remained. Altogether, we used two biological replicates (PADI2_mut1: 16,177 cells, PADI2mut2: 16,187 cells, WT1: 8,568 cells, WT2: 8,571 cells).

### Gene Ontology (GO) and Regulatory Gene Set analysis

GO Annotation and regulatory gene set analysis were performed using the online tool Gene Set Enrichment Analysis (GSEA, http://software.broadinstitute.org/gsea/index.jsp) collection database v5 [75,76]. The significant cut-off p-value and FDR q-value < 0.05. Plots were done with the use of Prism (GraphPad Prism 10.0.3 for MacOS), Cytoscape (version 3.9.1) and R4.0.2.

#### ChIP-qPCRs

For ChIP assays [30], 4 x 10^6^ of FACS sorted GFP-positive HeLa cells (WT-PADI2, L162A-PADI2, L162A/W161A-PADI2, L162A/W161A/T159A-PADI2) were cross-linked for 10 min with 1% formaldehyde at 37°C. The chromatin lysate was sonicated to a DNA fragment size range of 100-200bp using a Biorupter sonicator (Diagenode). CDK9 was immunoprecipitated with 15μg of an anti-CDK9 antibody (D-7, sc-13130, lot no # B1422) or control mouse IgG (12-371, Merck) in IP Buffer with 2X SDS buffer (100mM NaCl, 50mM Tris-HCl, pH8, 5mM EDTA and 0.5% SDS) and 1X Triton buffer (100mM Tris-HCl, pH8.8, 100mM NaCl, 5mM EDTA and 5% Triton-X) with protease inhibitors (11836170001, Roche) for 16 hours at 4° C. This step was followed by incubation with 50μl of Dynabeads® M-280 sheep anti-mouse IgG (11201D, Thermo Scientific) for 3 hours. Beads were washed 3 times with low salt buffer (140mM NaCl, 50mM HEPES, pH 7.4, 1% Triton-X 100), 2 times with high salt buffer (500 mM NaCl, 50mM HEPES, pH 7.4, 1% Triton-X 100) followed by single wash with LiCl Buffer (10mM Tris HCl pH 8.0, 250 mM LiCl, 1% NP-40, 1% sodium deoxycholic acid and 1mM EDTA) and 1× TE buffer at 4°C. Subsequently, crosslinks were reversed at 65° C overnight, followed by RNase treatment for 1.5 hours, and bound DNA was purified by Phenol-Chloroform extraction. The resulting eluted DNA was quantified by Qubit 3.0 Fluorometer (Life Technologies) and followed by real-time qPCR analysis. Data are represented as fold-change over input fraction from at least 3 biological replicate experiments. Primers used for qPCR are listed in **Table S1**.

### GFP-tagged PADI2 Immunoprecipitation, western blot

Briefly, 3 × 10^6^ FACS sorted GFP-positive (WT-PADI2, L162A-PADI2, L162A/W161A-PADI2, L162A/W161A/T159A-PADI2) cells were lysed on ice for 30 min in lysis buffer (1% Triton X-100 in 50mM Tris pH 7.4–7.6, 130 mM NaCl) containing protease inhibitors (11836170001, Roche) with rotation, followed by sonication for 7 min with every 30 sec on / 30 sec off. After centrifugation at 4°C and 13,000 rpm for 10 min, extracts were used for protein quantitation. For the immunoprecipitation (IP) assay, 2mg of extract was incubated for 12 hours with 100μl Dynabeads Protein A (10002D, Thermo Scientific). The monoclonal antibody anti-GFP (polyclonal rabbit, A-11122, Invitrogen) or a control rabbit IgG (2729S, Cell Signaling) was coupled with Dynabeads before incubation with extract at 4°C. The samples were washed 10 times with lysis buffer and boiled for 5 min in SDS gel sample buffer. Proteins were visualized by 4% to 12% SDS-PAGE gels and western blotting, using anti-GFP (11814460001, Roche), anti-CDK9 (sc-13130, Santa Cruz), or anti-CCNT1 (A303-499A, Bethyl Labs) were used for western blots.

### Multiple sequence alignments

All of the homolog PADI sequences available in the OMA database were downloaded [52]. The species for the study were selected using three criteria: (i) a good representation of the different mammalian lineages, (ii) good sequence quality, and (iii) the presence of a full PADI2 sequence. Three bird species (chicken, turkey, and duck) were used as outgroups. The protein sequences were aligned using the software PRANK [77] and a pruned mammalian guide tree with branch distances [78]. This program uses an evolutionary model to place insertions and deletions, minimizing the over-alignment of non-homologous regions, and has been shown to improve dN/dS estimates [79]. Local realignments of two regions were done using MAFFT [80] within AliView [81]. Another MSA using only one-to-one orthologous PADI2 sequences was also built. TrimAI [82] was applied to the PADI family MSA, enabling the -automated1 option, as recommended before a phylogenetic reconstruction. The PADI2 protein MSA was then converted into a nucleotide alignment using the script Pal2Nal. The corresponding cDNA sequences for each species were gathered from the OMA database and converted using the script Pal2Nal [83]. The MSAs generated and the trees used are available as Supplementary Material S1.

### Phylogenetic reconstruction

We reconstructed the PADI family tree using RaxML [84] with 200 bootstrap values (-N 200), with an optimal amino acid substitution model chosen automatically (-m PROTGAMMAAUTO), and a rapid bootstrap algorithm (-f a) along with reproducible seed values (-p 12345, -x 345). Two reconstructions were performed: one with the original PADI family MSA and another one with the trimmed version of the alignment.

### Estimation of dN/dS and positive selection

Estimates were made for both the number of non-synonymous substitutions per non-synonymous site (dN) and the number of synonymous substitutions per synonymous site (dS) using the free-ratio model in CodeML [85]. For each branch and leaf in the tree, we performed a branch-site test of positive selection [54], implemented in the Phylogenetic Analysis by Maximum Likelihood (PAML) software package [85] and available in the environment for tree exploration (ETE3) framework as bsA/bsA1 [86]. A likelihood ratio (LRT) was calculated as 2*(L1-L0), where L1 and L0 are the maximum likelihood value for the alternative hypothesis and the null hypothesis respectively. A chi-squared distribution with 1 degree of freedom was used to calculate the p-values. Only the positions for which the p-value (< 0.05) is significant for positive selection are reported.

### Characterization of positively selected residues using MVORFFIP

The prediction of interface residues was done using MVORFFIP [56]. MVORFFIP is a structure-based prediction method that identifies protein-, peptide-, RNA- and DNA interfaces based on a range of structural, evolutionary, experimental, and energy-based information integrated by a Random Forest classifier The structure of PADI2 was submitted to the MVORFFIP server (http://www.bioinsilico.org/MVORFFIP) and protein interface scores were assigned to individual amino acids.

### Structural modeling of PADI2-CDK9/CCNT1 and selection of putative interface residues

The structure of the trimer complex, PADI2-CDK9, was derived using protein docking as follows. The crystal structure of PADI2 (PDB code: 4n2a) [25] and CDK9/CCNT1 (PDB code: 3blr) [59] were available. The protein docking was performed using VD^2^OCK [87], generating the ensemble of over 10000 docking poses. Docking conformers were clustered using the GROMACS [88] and sorted by the ES3DC potential [89]. The top 200 poses were selected to identify the top 10 putative interface residues: for this, a composite score was calculated using the docking information, the MVORFFIP score and sequence conservation. Among the top 200 docking poses, as discussed above, a normalized score (Z score) was calculated for each exposed residue in PADI2. The mean and standard deviations to calculate the Z score were computed over the entire docking space, i.e., considering all docking poses. The MVORFFIP score was also computed as shown above. Finally, the conservation of each residue was computed using al2co [90].

### AlphaFold-Multimer modeling of PADI2-CDK9/CCNT1 complex

Structural models of the trimer PADI2-CDK9/CCNT1 were generated using AlphaFold-Multimer [60] as follows. The sequences were retrieved as mentioned above, PADI2 (PDB code: 4n2a) [25] and CDK9/CCNT1 (PDB code: 3blr) [59]. The structure was derived using model parameters corresponding to “model_1_multimer”.

## Data Availability

The OMA is an open-source database for the inference of orthologs among complete genomes https://omabrowser.org/oma/home/. ETE3 was used for the reconstruction, analysis, and visualization of phylogenetic trees and is available at http://etetoolkit.org. Other open-source programs used here were GROMACS, VD2OCK, MVORFFIP, and al2co, which are available at http://www.gromacs.org, http://www.bioinsilico.org/VD2OCK, http://www.bioinsilico.org/MVORFFIP, and http://prodata.swmed.edu/download/pub/al2co/, respectively. File S2.pse, File S3.pse, File S4.pse, and File S5.pse correspond to PyMOL (http://pymol.org) sessions created to produce Supplementary Movie S1.mpg, Supplementary Movie S2.mpg, Supplementary Movie S3.mpg, and Movie S4.mpg respectively. The PyMOL sessions also give access to the coordinates of the PADI2 and structural model of the PADI2/CDK9-CCNT1 complex. Single-cell RNA sequencing data performed and used in this study have been deposited at GEO under the accession code GSE246420.

## Acknowledgment

We thank Fátima Gebauer and François Le Dily from CRG for their constructive criticism and advice on this manuscript. We acknowledge Veronica A. Raker for manuscript editing. We thank the CNAG (National Center for Genomic Analysis, Barcelona) Genomic facility and CRG flow Cytometry unit for all technical support.

## Author Contributions

Conceptualization, P.S.; Methodology, J.L.V., N.F.F., P.S.; Investigation, J.L.V., N.F.F., B.O., M.B., and P.S.; Formal Analysis, J.L.V., N.F.F., and P.S.; Data Curation, J.L.V., N.F.F., M.E.R., C.T., F.P.P., B.O., C.N., P.S.; Project Administration, P.S., Writing-Original Draft, P.S. Writing-Review & Editing, J.L.V., N.F.F., B.O., M.E.R., C.T., F.P.P., M.B., C.N., P.S.; Funding Acquisition, P.S.; Supervision, P.S.

## Supplementary Data

Supplementary data are available online.

