## Supplementary text Figures and tables for "Evolutionary analysis reveals the role of a non-catalytic domain of peptidyl arginine deiminase 2 in transcriptional regulation": JLL and NFF Sup combined 2023_compressed.pdf

**This PDF file includes:**

Supplementary text
Figs S1 to S5
Tables S1- S4
Legends for movies S1 to S4
Legends for files S1 to S5

**Other supplementary materials for this manuscript include the following:**

Movies S1 to S4
Files S1 to S5

**Supplementary figures and table legends**

**S1 Fig (related to Fig 1).** Topology tree showing the distribution of species**.** Each node is numbered for easier identification of branches under positive selection.

**S2 Fig (related to Fig 1).** Phylogenetic tree of the PADI family of enzymes reconstructed using RaxML. The five different PADI genes are purple: *PADI1* (green), *PADI2* (purple), *PADI3* (blue), *PADI4* (red), and *PADI6* (orange). Branch lengths represent the phylogenetic distance between sequences.

**S3 Fig (related to Fig 1).** Phylogenetic reconstruction of all PADIs. Branches are colored according to the bootstrap values; blue: ≥75, green: ≥50.

**S4 Fig (related to Fig 1, Fig 4, Files S3-S4, and Movies S2-S3)**. (**A**-**C**). Gene order and synteny for all identified PADI genes in three representative species: duck (top), human (middle), and mouse (bottom). Gene locations were obtained from Ensembl v97. (**D-E** Structural model of PADI2 interaction with the P-TEFb complex. Structural model of PADI2-CDK9/CCNT1 complex derived by: **(D)** Docking, showing one of the top-ranked docking poses. (**E)** Using AlphaFold-Multimer; In both panels, PADI2 is shown as a cartoon using the same color scheme as in Figure 3, and CDK9 and CCNT1 are shown in cyan and magenta, respectively; L162 is shown in blue using sphere representation. The PyMol session of PADI2 with the P-TEFb complex with docking and AlphaFold-Multimer is available in Supplementary Material File S3 and S4, respectively**.**

**S5 Fig (related to Fig 5)**. HeLa cells expressing the GFP-tagged PADI2 were used for GFP-positive FACS. (**A**). Schematics outlining the experimental strategy. HeLa cells were transfected with indicated GFP-tagged PADI2 plasmids, followed by sorting of GFP-positive cells. The gating strategy for isolating GFP-expressing cells using the BD Influx cell sorter is shown. Top panel, is a strategy used for control cells; the bottom panel, is a strategy used for GFP-expressing cells. (A1, B1) A SSC-H (side scatter height) versus FSC-H (forward scatter height), morphological-related parameter dot-plot was used to exclude debris by gating cells in G1. (A2, B2) Doublets were excluded using FSC-H versus FSC-A (forward scatter area) dot-plot by gating singlets in G2. (A3, B3) Dead cells were excluded using DAPI versus FSC-A dot-plot by gating living cells in G3. (A4, B4) GFP-positive cells. (**B**)**.** Representative microscopy images of HeLa cells expressing PADI2 as the control (–GFP) or a GFP-tagged wild-type (WT), single mutant (T159A, W161A or L162A), double mutant (L162A/W161A), or triple mutant (L162A/W161A/T159A) form. (**C**). The growth curve displaying percentage confluence was determined using the Incucyte® live cell analysis system over time for HeLa cells specifically expressing the WT PAD12 or mutant PAD12 (single, double, or triple, as for (B). Data represent mean ± SD. ***p value < 0.001.

**S6 Fig (related to Fig 6)**. (**A**). A t-distributed stochastic neighbor embedding plot (tSNE) depicting unsupervised clustering leads to three different populations that appear to be descriptionally distinct. (**B**). An overlap of (L162A/W161A/T159A) mutant vs. WT PADI2 tSNE shows that these different clusters are not directly associated with the mutational status of PADI2. (**C**). Hypothetical evolutionary model of positively selected residues to coordinate with the circumferential residues to facilitate protein–protein interactions to contribute to cellular functions.

**Table S1- Primers used in this study.**

**RT-qPCR**

| Name | Sequence (5′ to 3′) |
|---|---|
| *CENPQ, F.P.* | CAACCTCTGTCAAAGAGTACC |
| *CENPQ, R.P.* | TCAGGAAGTTGAGATGGTATTG |
| *c-MYC*, *F.P.* | AGAGTCTGGATCACCTTCTG |
| *c-MYC*, *R.P* | GGTTGTTGCTGATCTGTCTC |
| *CCNB1, F.P.* | GGTAACAAAGTCAGTGAACAAC |
| *CCNB, R.P* | TGGCACCAGCATAGGTA |
| GAPDH, *F.P.* | GAGTCAACGGATTTGGTCGT |
| GAPDH, *R.P* | TTGATTTTGGAGGGATCTCG |
| *LRRC39, F.P.* | CCCTTCTTCTCTGCTGAAAC |
| *LRRC39, R.P* | CTGAGAATCAGTTCCTGAAGTC |
| *PADI2 F.P.* | CATGTCCCAGATGATCCTGC |
| *PADI2 R.P* | CATGGTAGAGCTTCCGCC |
| | **ChIP-qPCR** |
| *CENPQ*-TSS, *F.P.* | GGTCTCCTCTGTTTACGAATG |
| *CENPQ*-TSS, *R.P.* | GCATTTGCTTTACCAGACATC |
| *c-MYC*-TSS *F.P.* | CTGCTTAGACGCTGGATTT |
| *c-MYC*-TSS *R.P* | TTTAGGCATTCGACTCATCTC |
| *CCNB1*-TSS *F.P.* | CCTGGAAACGCATTCTCTG |
| *CCND2*-TSS *R.P* | cCAGCCTAGCCTCAGATTTA |
| *LRRC39* TSS *F.P.* | GGTCCTTTCTGAAATGGTATC |
| *LRRC39* TSS *R.P* | TCAGGTCTTCATTGAGTTTC |

**Table S2** – Differentially expressed genes in PADI2 triple mutant (L162A/W161A/T159A) versus PADI2 WT as indicated in **Fig 6A-B.**

**Table S3** – List of significantly enriched molecular functions and biological processes in PADI2 triple mutant (L162A/W161A/T159A) /PADI2 WT related to **Fig 6C.**

**Table S4-** Set of gene regulatory elements in upregulated and downregulated genes as shown in **Fig 6 D-E.**

**Supplementary material files and movie legends**

**Material file S1.** Multiple sequence alignment files were generated in amino acids and cDNA, as well as the reconstructed tree in Newick format.

**Material file S2.** PyMOL session file of PADI2 (green), with the CTD-R1810 peptide complex highlighting the positively selected amino acid residues.

**Material file S3.** PyMOL session file of docking model of PADI2 (green), with CDK9/CCNT1 (in cyan and magenta, respectively).

**Material file S4.** PyMOL session file of AlphaFold-Multimer model of PADI2 (green), with CDK9/CCNT1 (in cyan and magenta, respectively).

**Material file S5.** PyMOL session file of AlphaFold-Multimer & docking shared region of PADI2 (green) centered around L162 encompassing loop with CDK9/CCNT1 (in cyan and magenta, respectively).

**Movie S1-** A movie showing a 360-degree rotation of the lowest energy model of PADI2-with R1810 RNAP2 peptide complex. All 20 positively selected residues are highlighted as shown in Figure 3.

**Movie S2-** A movie depicting a 360-degree rotation docking model of PADI2-CDK9/CCNT1 complex, with PADI2 in green shades, CDK9 in cyan, and CCNT1 in magenta. L162 is depicted in dark blue with a sphere representation.

**Movie S3-** A movie showing a 360-degree rotation of AlphaFold-Multimer structural model of PADI2-CDK9/CCNT1 complex, with PADI2 in green shades, CDK9 in pale cyan, and CCNT1 in pink. L162 is depicted in dark blue with a sphere representation.

**Movie S4-** A movie illustrating a 360-degree rotation of PADI2 in green with the shared interface between docking and AlphaFold-Multimer models in red and the center of mass of the top 200 docking poses represented as red spheres. T159, L162, and W161 are shown in sphere representations, with L162 depicted in dark blue, and T159 and W161 in cyan.

### S1 Fig

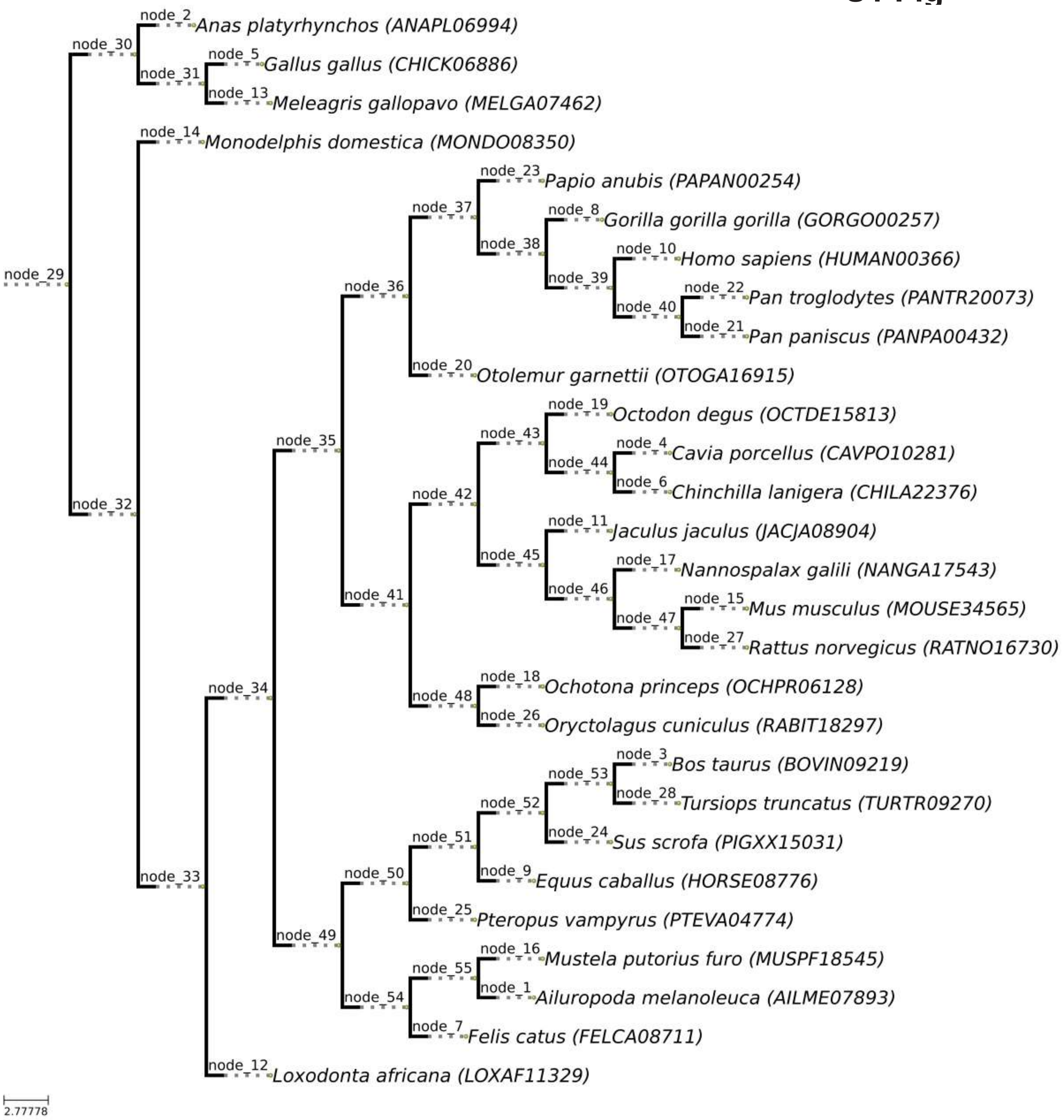
node_29
node_30
node_2
Anas platyrhynchos (ANAPL06994)
node_31
node_5
Gallus gallus (CHICK06886)
node_13
Meleagris gallopavo (MELGA07462)
node_32
node_14
Monodelphis domestica (MONDO08350)
node_33
node_34
node_35
node_36
node_37
node_23
Papio anubis (PAPAN00254)
node_38
node_8
Gorilla gorilla gorilla (GORGO00257)
node_39
node_10
Homo sapiens (HUMAN00366)
node_40
node_22
Pan troglodytes (PANTR20073)
node_21
Pan paniscus (PANPA00432)
node_20
Otolemur garnettii (OTOGA16915)
node_41
node_42
node_43
node_19
Octodon degus (OCTDE15813)
node_44
node_4
Cavia porcellus (CAVPO10281)
node_6
Chinchilla lanigera (CHILA22376)
node_45
node_11
Jaculus jaculus (JACJA08904)
node_46
node_17
Nannospalax galili (NANGA17543)
node_47
node_15
Mus musculus (MOUSE34565)
node_27
Rattus norvegicus (RATNO16730)
node_48
node_18
Ochotona princeps (OCHPR06128)
node_26
Oryctolagus cuniculus (RABIT18297)
node_49
node_50
node_51
node_52
node_53
node_3
Bos taurus (BOVIN09219)
node_28
Tursiops truncatus (TURTR09270)
node_24
Sus scrofa (PIGXX15031)
node_9
Equus caballus (HORSE08776)
node_25
Pteropus vampyrus (PTEVA04774)
node_54
node_55
node_16
Mustela putorius furo (MUSPF18545)
node_1
Ailuropoda melanoleuca (AILME07893)
node_7
Felis catus (FELCA08711)
node_12
Loxodonta africana (LOXAF11329)
2.77778

S2 Fig

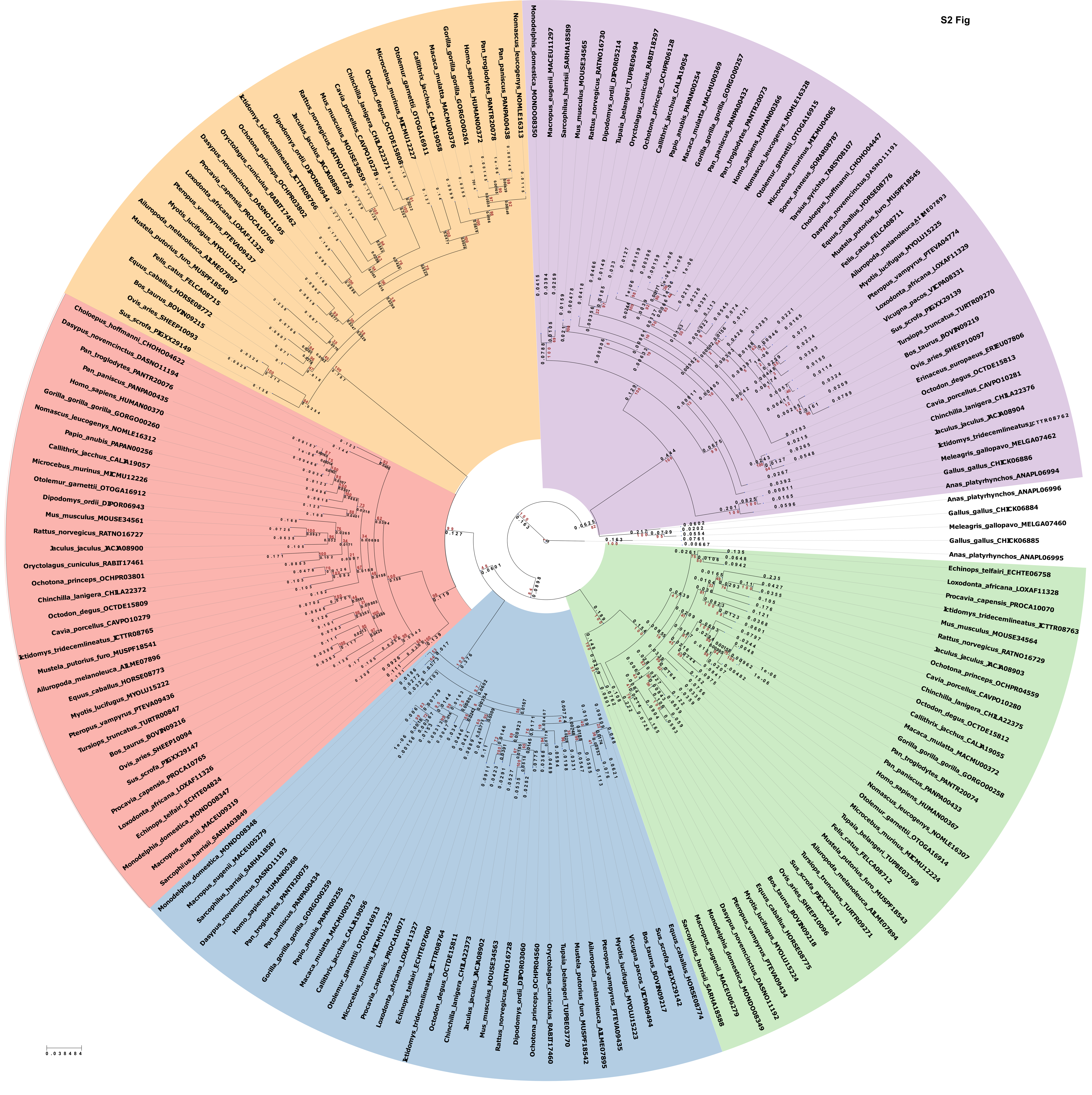
Nomascus_leucogenys_NOMLE16313
Pan_paniscus_PANPA00438
Pan_troglodytes_PANTR20078
Homo_sapiens_HUMAN00372
Gorilla_gorilla_gorilla_GORGO00261
Macaca_mulatta_MACMU00376
Callithrix_jacchus_CALJA19058
Otolemur_garnettii_OTOGA16911
Microcebus_murinus_MICMU12227
Octodon_degus_OCTDE15808
Chinchilla_lanigera_CHILA22371
Cavia_porcellus_CAVPO10278
Mus_musculus_MOUSE34559
Rattus_norvegicus_RATNO16726
Jaculus_jaculus_JACJA08899
Dipodomys_ordii_DIPOR06944
Ictidomys_tridecemlineatus_ICTTR08766
Ochotona_princeps_OCHPR03802
Oryctolagus_cuniculus_RABIT17462
Dasypus_novemcinctus_DASNO11195
Procavia_capensis_PROCA10766
Loxodonta_africana_LOXAF11325
Pteropus_vampyrus_PTEVA09437
Myotis_lucifugus_MYOLU15221
Ailuropoda_melanoleuca_AILME07897
Mustela_putorius_furo_MUSPF18540
Felis_catus_FELCA08715
Equus_caballus_HORSE08772
Bos_taurus_BOVIN09215
Ovis_aries_SHEEP10093
Sus_scrofa_PIGXX29149
Choloepus_hoffmanni_CHOHO04622
Dasypus_novemcinctus_DASNO11194
Pan_troglodytes_PANTR20076
Pan_paniscus_PANPA00435
Homo_sapiens_HUMAN00370
Gorilla_gorilla_gorilla_GORGO00260
Nomascus_leucogenys_NOMLE16312
Papio_anubis_PAPAN00256
Callithrix_jacchus_CALJA19057
Microcebus_murinus_MICMU12226
Otolemur_garnettii_OTOGA16912
Dipodomys_ordii_DIPOR06943
Mus_musculus_MOUSE34561
Rattus_norvegicus_RATNO16727
Jaculus_jaculus_JACJA08900
Oryctolagus_cuniculus_RABIT17461
Ochotona_princeps_OCHPR03801
Chinchilla_lanigera_CHILA22372
Octodon_degus_OCTDE15809
Cavia_porcellus_CAVPO10279
Ictidomys_tridecemlineatus_ICTTR08765
Mustela_putorius_furo_MUSPF18541
Ailuropoda_melanoleuca_AILME07896
Equus_caballus_HORSE08773
Myotis_lucifugus_MYOLU15222
Pteropus_vampyrus_PTEVA09436
Tursiops_truncatus_TURTR00847
Bos_taurus_BOVIN09216
Ovis_aries_SHEEP10094
Sus_scrofa_PIGXX29147
Procavia_capensis_PROCA10765
Loxodonta_africana_LOXAF11326
Echinops_telfairi_ECHTE04824
Monodelphis_domestica_MONDO08347
Macropus_eugenii_MACEU09319
Sarcophilus_harrisii_SARHA03849
Monodelphis_domestica_MONDO08348
Macropus_eugenii_MACEU05279
Sarcophilus_harrisii_SARHA18587
Dasypus_novemcinctus_DASNO11193
Homo_sapiens_HUMAN00368
Pan_troglodytes_PANTR20075
Pan_paniscus_PANPA00434
Gorilla_gorilla_gorilla_GORGO00259
Papio_anubis_PAPAN00255
Macaca_mulatta_MACMU00373
Callithrix_jacchus_CALJA19056
Otolemur_garnettii_OTOGA16913
Microcebus_murinus_MICMU12225
Procavia_capensis_PROCA10071
Loxodonta_africana_LOXAF11327
Echinops_telfairi_ECHTE07600
Ictidomys_tridecemlineatus_ICTTR09764
Octodon_degus_OCTDE15811
Chinchilla_lanigera_CHILA22373
Jaculus_jaculus_JACJA08902
Mus_musculus_MOUSE34563
Rattus_norvegicus_RATNO16728
Dipodomys_ordii_DIPOR03060
Ochotona_princeps_OCHPR04560
Oryctolagus_cuniculus_RABIT17460
Tupaia_belangeri_TUPBE03770
Mustela_putorius_furo_MUSPF18542
Ailuropoda_melanoleuca_AILME07895
Pteropus_vampyrus_PTEVA09435
Myotis_lucifugus_MYOLU15223
Vicugna_pacos_VICPA09494
Bos_taurus_BOVIN09217
Sus_scrofa_PIGXX29142
Equus_caballus_HORSE08774
Sarcophilus_harrisii_SARHA18588
Macropus_eugenii_MACEU06279
Monodelphis_domestica_MONDO08349
Dasypus_novemcinctus_DASNO11192
Pteropus_vampyrus_PTEVA09434
Myotis_lucifugus_MYOLU15224
Equus_caballus_HORSE08775
Bos_taurus_BOVIN09218
Ovis_aries_SHEEP10096
Sus_scrofa_PIGXX29141
Tursiops_truncatus_TURTR09271
Ailuropoda_melanoleuca_AILME07894
Mustela_putorius_furo_MUSPF18543
Felis_catus_FELCA08712
Tupaia_belangeri_TUPBE03769
Microcebus_murinus_MICMU12224
Otolemur_garnettii_OTOGA16914
Nomascus_leucogenys_NOMLE16307
Homo_sapiens_HUMAN00367
Pan_paniscus_PANPA00433
Pan_troglodytes_PANTR20074
Gorilla_gorilla_gorilla_GORGO00258
Macaca_mulatta_MACMU00372
Callithrix_jacchus_CALJA19055
Octodon_degus_OCTDE15812
Chinchilla_lanigera_CHILA22375
Cavia_porcellus_CAVPO10280
Ochotona_princeps_OCHPR04559
Jaculus_jaculus_JACJA08903
Rattus_norvegicus_RATNO16729
Mus_musculus_MOUSE34564
Ictidomys_tridecemlineatus_ICTTR08763
Procavia_capensis_PROCA10070
Loxodonta_africana_LOXAF11328
Echinops_telfairi_ECHTE06758
Anas_platyrhynchos_ANAPL06995
Gallus_gallus_CHICK06885
Meleagris_gallopavo_MELGA07460
Gallus_gallus_CHICK06884
Anas_platyrhynchos_ANAPL06996
Anas_platyrhynchos_ANAPL06994
Gallus_gallus_CHICK06886
Meleagris_gallopavo_MELGA07462
Ictidomys_tridecemlineatus_ICTTR08762
Jaculus_jaculus_JACJA08904
Chinchilla_lanigera_CHILA22376
Cavia_porcellus_CAVPO10281
Octodon_degus_OCTDE15813
Erinaceus_europaeus_ERIEU07806
Ovis_aries_SHEEP10097
Bos_taurus_BOVIN09219
Tursiops_truncatus_TURTR09270
Sus_scrofa_PIGXX29139
Vicugna_pacos_VICPA08331
Loxodonta_africana_LOXAF11329
Pteropus_vampyrus_PTEVA04774
Myotis_lucifugus_MYOLU15225
Ailuropoda_melanoleuca_AILME07893
Felis_catus_FELCA08711
Mustela_putorius_furo_MUSPF18545
Equus_caballus_HORSE08776
Dasypus_novemcinctus_DASNO11191
Choloepus_hoffmanni_CHOHO04447
Tarsius_syrichta_TARSY08107
Sorex_araneus_SORAR08787
Microcebus_murinus_MICMU04065
Otolemur_garnettii_OTOGA16915
Nomascus_leucogenys_NOMLE16328
Homo_sapiens_HUMAN00366
Pan_troglodytes_PANTR20073
Pan_paniscus_PANPA00432
Gorilla_gorilla_gorilla_GORGO00257
Macaca_mulatta_MACMU00369
Papio_anubis_PAPAN00254
Callithrix_jacchus_CALJA19054
Ochotona_princeps_OCHPR06128
Oryctolagus_cuniculus_RABIT18297
Tupaia_belangeri_TUPBE09494
Dipodomys_ordii_DIPOR05214
Rattus_norvegicus_RATNO16730
Mus_musculus_MOUSE34565
Sarcophilus_harrisii_SARHA18589
Macropus_eugenii_MACEU11297
Monodelphis_domestica_MONDO08350
0.038484

S3 Fig

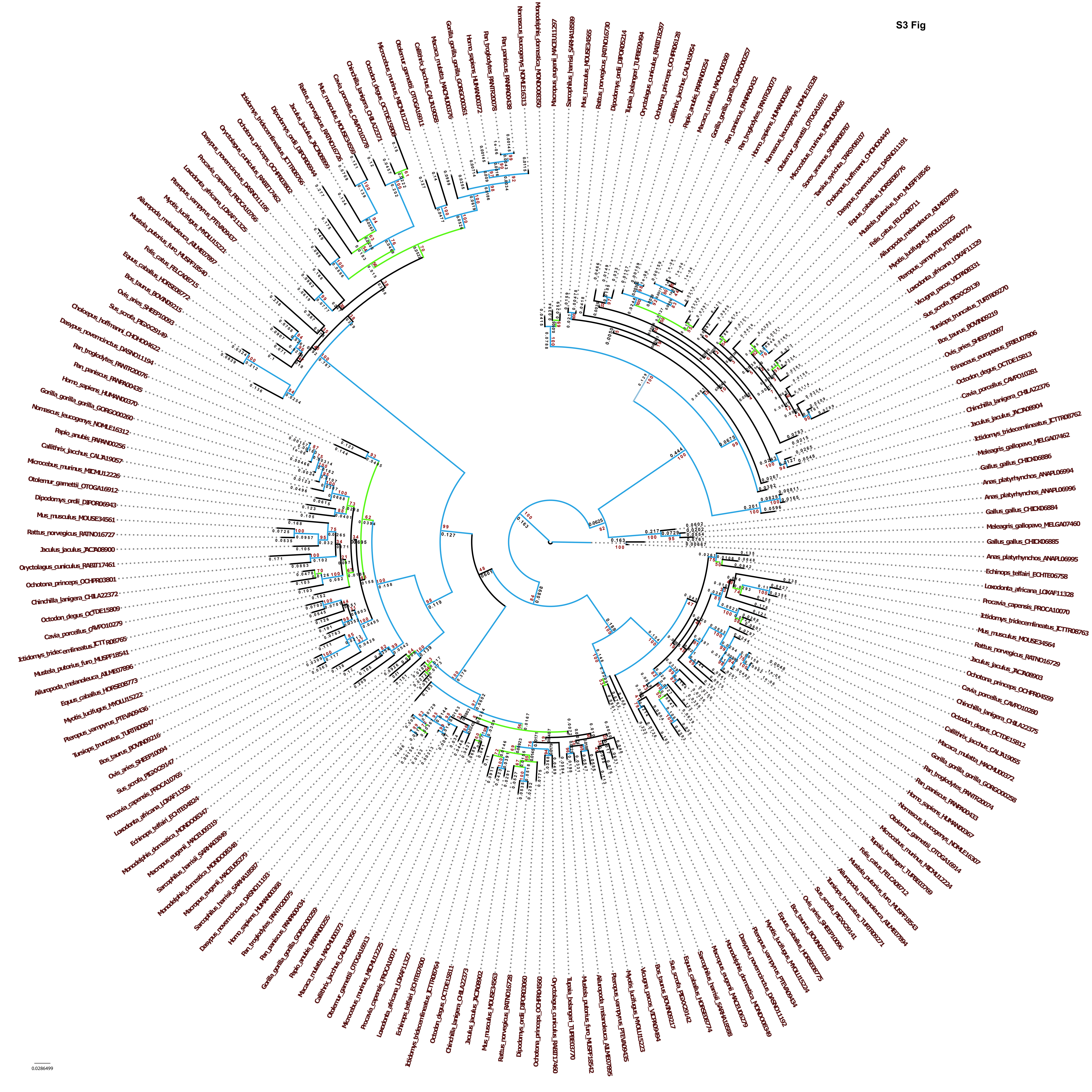

S4 Fig

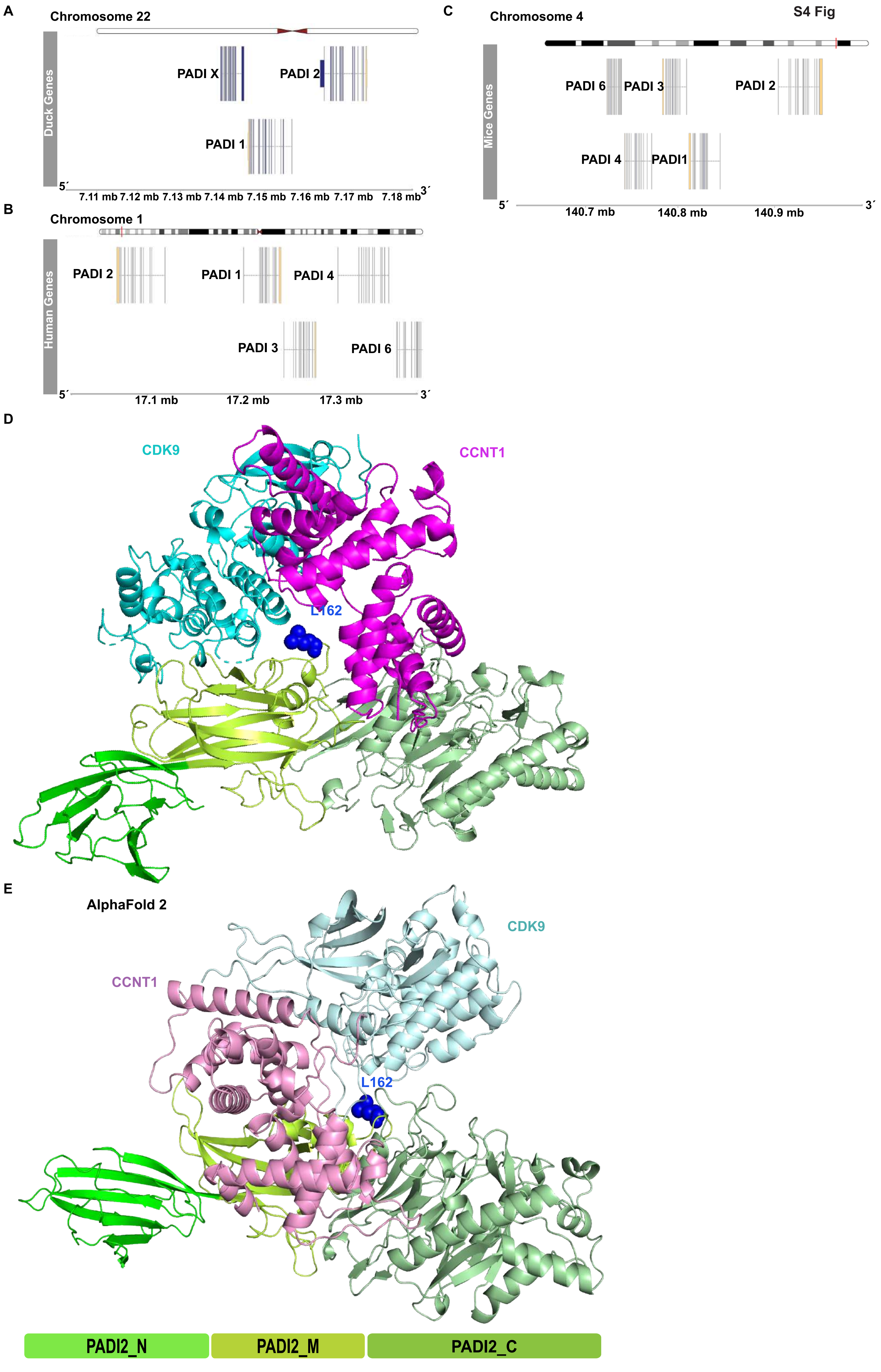
A
Chromosome 22
Duck Genes
PADI X
PADI 2
PADI 1
5´
3´
7.11 mb 7.12 mb 7.13 mb 7.14 mb 7.15 mb 7.16 mb 7.17 mb 7.18 mb
B
Chromosome 1
Human Genes
PADI 2
PADI 1
PADI 4
PADI 3
PADI 6
5´
3´
17.1 mb
17.2 mb
17.3 mb
C
Chromosome 4
Mice Genes
PADI 6
PADI 3
PADI 2
PADI 4
PADI1
5´
3´
140.7 mb
140.8 mb
140.9 mb
D
CDK9
CCNT1
L162
E
AlphaFold 2
CDK9
CCNT1
L162
PADI2_N
PADI2_M
PADI2_C

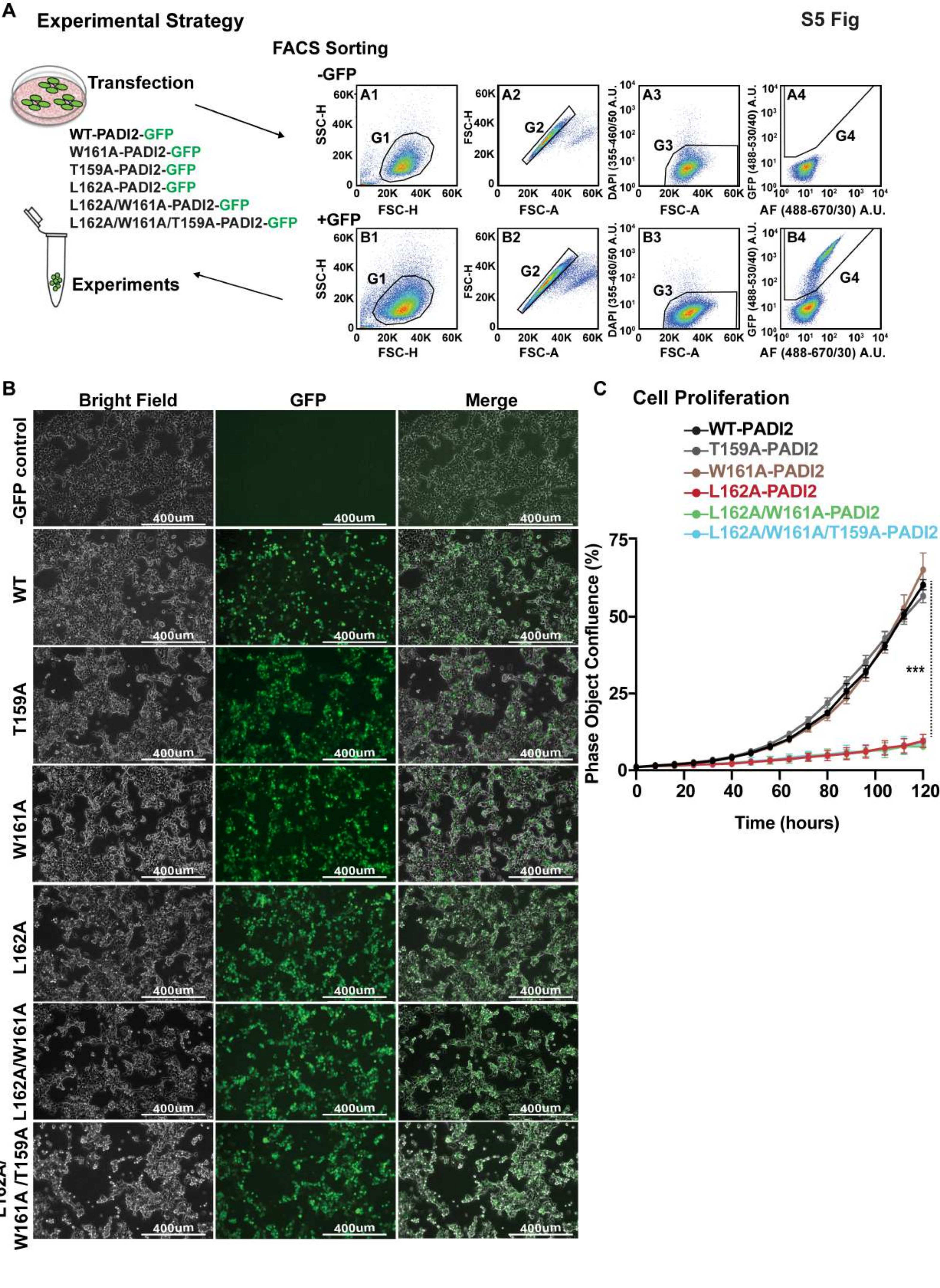
S5 Fig
A
Experimental Strategy
Transfection
WT-PADI2-GFP
W161A-PADI2-GFP
T159A-PADI2-GFP
L162A-PADI2-GFP
L162A/W161A-PADI2-GFP
L162A/W161A/T159A-PADI2-GFP
Experiments
FACS Sorting
-GFP
+GFP
A1
A2
A3
A4
B1
B2
B3
B4
G1
G2
G3
G4
SSC-H
FSC-H
FSC-A
DAPI (355-460/50 A.U.
GFP (488-530/40) A.U.
AF (488-670/30) A.U.
B
Bright Field
GFP
Merge
-GFP control
WT
T159A
W161A
L162A
L162A/W161A
L162A/W161A /T159A
400um
C
Cell Proliferation
WT-PADI2
T159A-PADI2
W161A-PADI2
L162A-PADI2
L162A/W161A-PADI2
L162A/W161A/T159A-PADI2
Phase Object Confluence (%)
Time (hours)
***

S6 Fig

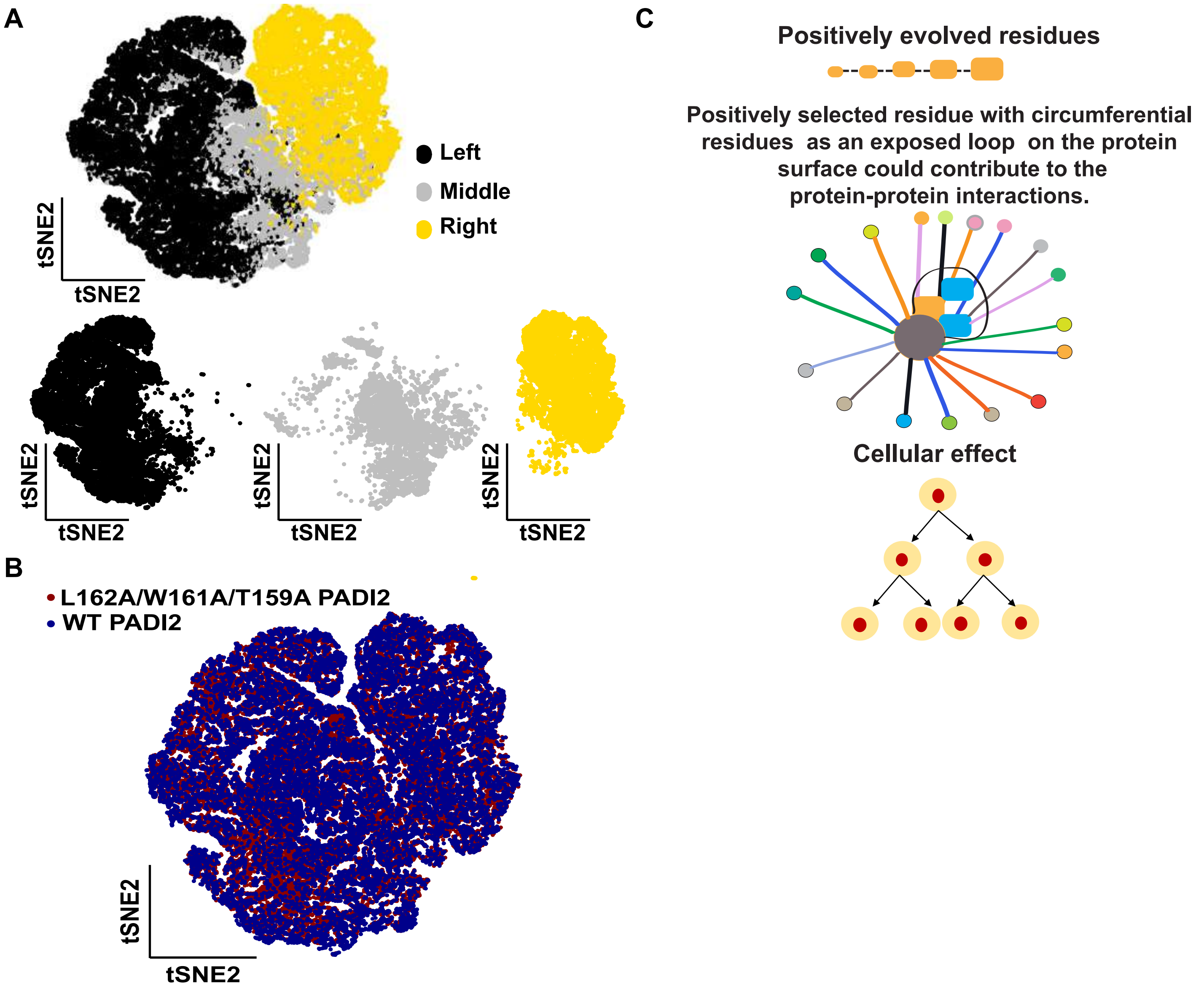
A
Left
Middle
Right
tSNE2
tSNE2
B
L162A/W161A/T159A PADI2
WT PADI2
tSNE2
tSNE2
C
Positively evolved residues
Positively selected residue with circumferential residues as an exposed loop on the protein surface could contribute to the protein-protein interactions.
Cellular effect
